# An mRNA processing pathway suppresses metastasis by governing translational control from the nucleus

**DOI:** 10.1101/2021.10.04.463118

**Authors:** Albertas Navickas, Hosseinali Asgharian, Juliane Winkler, Lisa Fish, Kristle Garcia, Daniel Markett, Martin Dodel, Bruce Culbertson, Sohit Miglani, Tanvi Joshi, Phi Nguyen, Steven Zhang, Nicholas Stevers, Hun-Way Hwang, Faraz Mardakheh, Andrei Goga, Hani Goodarzi

**Affiliations:** Department of Biochemistry and Biophysics, University of California, San Francisco, San Francisco, CA, USA; Department of Urology, University of California, San Francisco, San Francisco, CA, USA; Helen Diller Family Comprehensive Cancer Center, University of California, San Francisco, San Francisco, CA, USA; Bakar Computational Health Sciences Institute, University of California, San Francisco, San Francisco, CA, USA; Department of Cell and Tissue Biology, University of California, San Francisco, San Francisco, CA, USA; Centre for Cancer Cell and Molecular Biology, Barts Cancer Institute, Queen Mary University of London, London, UK; Department of Pathology, University of Pittsburgh, Pittsburgh, PA, USA; Department of Medicine, University of California, San Francisco, San Francisco, CA, USA

## Abstract

Cancer cells often co-opt post-transcriptional regulatory mechanisms to achieve pathologic expression of gene networks that drive metastasis. Translational control is a major regulatory hub in oncogenesis, however its effects on cancer progression remain poorly understood. To address this, we used ribosome profiling to compare genome-wide translation efficiencies of poorly and highly metastatic breast cancer cells and patient-derived xenografts. We developed novel regression-based methods to analyze ribosome profiling and alternative polyadenylation data, and identified HNRNPC as a translational controller of a specific mRNA regulon. Mechanistically, HNRNPC, in concert with PABPC4, binds near to poly(A) signals, thereby governing the alternative polyadenylation of a set of mRNAs. We found that HNRNPC and PABPC4 are downregulated in highly metastatic cells, which causes HNRNPC-bound mRNAs to undergo 3’ UTR lengthening and subsequently, translational repression. We showed that modulating HNRNPC expression impacts the metastatic capacity of breast cancer cells in xenograft mouse models. We also found that a small molecule, previously shown to induce a distal-to-proximal poly(A) site switching, counteracts the HNRNPC-PABPC4 driven deregulation of alternative polyadenylation and decreases the metastatic lung colonization by breast cancer cells *in vivo*.

## INTRODUCTION

Metastasis is the leading cause of cancer-related mortality^1^, and understanding its molecular underpinnings remains a challenge in basic and translational biology. Although genetic alterations can contribute to cancer progression^2^, the majority of known cellular routes to metastasis involve non-genetic mechanisms^1^. Cancer cells often co-opt post-transcriptional regulatory networks to activate pro-metastatic gene expression programs^3–5^. Therefore, all stages of the mRNA life cycle—including alternative splicing, post-transcriptional modification, translation, and decay—have been implicated in cancer progression^4–6^. Translational control has been increasingly recognized as an important regulatory node in tumorigenesis^7^, however our understanding on how translational deregulation acts in the later stages of cancer remains incomplete.

Translational control is tightly intertwined with other aspects of RNA biology, including the tRNA availability, the interplay between alternative 5’ and 3’ untranslated regions (UTRs), and their interaction with RNA binding proteins and microRNAs (miRNAs). To activate one of the main routes to metastasis, cancer cells have been shown to exploit the translational upregulation of several factors involved in the epithelial-to-mesenchymal transition^8,9^. Furthermore, numerous studies have observed a global tendency towards 3’ UTR shortening in cancer^10–14^, suggesting consequences from reduced interactions with RNA binding proteins and miRNAs^15^, including altered translation^7^. In some cases, these observations could be attributed to changes in expression of specific mRNA cleavage and polyadenylation factors^12,13^, although in many instances the underlying molecular mechanisms remain unknown. Similarly, we have previously demonstrated that the translational reprogramming that accompanies changes in tRNA expression landscape drives metastasis in breast cancer^16^. Importantly, a systematic characterization of translational control and its links to other aspects of RNA metabolism in metastasis is still lacking.

Here, we applied genome-wide experimental and computational approaches to address the changes in mRNA translation that accompany the metastatic progression in breast cancer. We performed ribosome profiling in both cell line- and patient-derived models of breast cancer metastasis, and used Ribolog, a novel analytical framework, to identify the underlying regulatory programs that govern changes in the translational control landscape. By applying these tools, we identified a functional interplay between nuclear RNA processing and translational control that disrupts the expression of a metastasis-suppressive regulon. Mechanistically, we found that the downregulation of HNRNPC and its interacting partner PABPC4 in highly metastatic cells was associated with 3’ UTR lengthening of their target mRNAs, resulting in their translational repression in part through an Argonaute-dependent pathway. HNRNPC depletion increased the metastatic capacity of breast cancer cells in xenograft models of metastasis and HNRNPC expression was negatively associated with clinical outcomes in multiple breast cancer patient datasets. We specifically identified PDLIM5, a cytoskeleton-associated protein implicated in mechanosensing, as a target of the HNRNPC-PABPC4 regulatory function that acts as a metastasis suppressor. We also showed that counteracting the 3’ UTR lengthening in highly metastatic breast cancer cells using a small molecule drug diminished their metastatic capacity *in vivo*, establishing this pathway as a potential therapeutic vulnerability.

## RESULTS

### Translational reprogramming accompanies metastatic progression in breast cancer

To capture changes in the translational landscape that are associated with breast cancer metastasis we performed ribosome profiling (Ribo-seq) on a commonly used triple receptor negative model of breast cancer metastasis, MDA-MB-231 breast cancer cells and their lung metastatic derivative cell line, MDA-LM2^17^. We predominantly recovered 33-34 nucleotide long ribosome protected mRNA footprints, aligning in frame with annotated coding sequences^18^, confirming the high quality of our dataset (Fig. S1a-b). We then sought to measure relative changes in translational activity genome-wide by calculating translational efficiency ratios (TER) between MDA-LM2 and parental MDA-MB-231 cells and identifying genes that are significantly up or down-regulated at the translational level.

To perform reliable differential analysis of Ribo-seq data and systematically account for possible confounders, we developed a new analytical framework for comparison of translation efficiencies (TE, representing the ratio between ribosome protected mRNA footprint and mRNA abundance sequencing read counts), aiming for as few *a priori* assumptions as possible (see Methods for details). The resulting method, which we have named Ribolog, relies on logistic regression to model individual Ribo-seq and RNA-seq reads in order to estimate logTER (i.e. log fold-change in TE) and its associated *p*-value across the coding transcriptome. Ribolog offers several advantages over the existing methods: (*i*) it does not assume a negative binomial (NB) distribution of read counts, and thus does not require estimation of the NB dispersion parameter; (*ii*) it detects and eliminates the translation stalling bias before estimating logTER; and (*iii*) it introduces the ribosome profiling-specific quality control metrics.

First, we used Ribolog to calculate the translation efficiency changes between poorly and highly metastatic breast cancer cells, and detected numerous differentially translated mRNAs (Fig. 1a). We then assessed the impact of changes in translation efficiency on the proteome by comparing the abundance of proteins in MDA-LM2 and parental MDA-MB-231 cells using tandem mass tag labeling and mass spectrometry (TMT-MS). As expected, we observed broad changes in the proteome as cells become more metastatic (Fig. S1c). Moreover, we observed that the changes in protein levels can be partially but not completely explained by changes in the mRNA levels (R = 0.5, *p* < 2 × 10^−16^ between mRNA and protein log fold-changes), which points to regulators of protein synthesis and decay as another source of variations. To formalize the changes in translation efficiency as a key factor in the observed modulations, we corrected changes in the protein levels by their respective changes in mRNA abundance, and observed that the resulting measure is significantly correlated with logTER values from Ribo-seq (R = 0.3, *p* < 2 × 10^−16^; Fig. S1d). These findings suggest that post-transcriptional regulation of translation efficiency has a significant impact on protein levels in highly metastatic cells.

**Figure 1.**
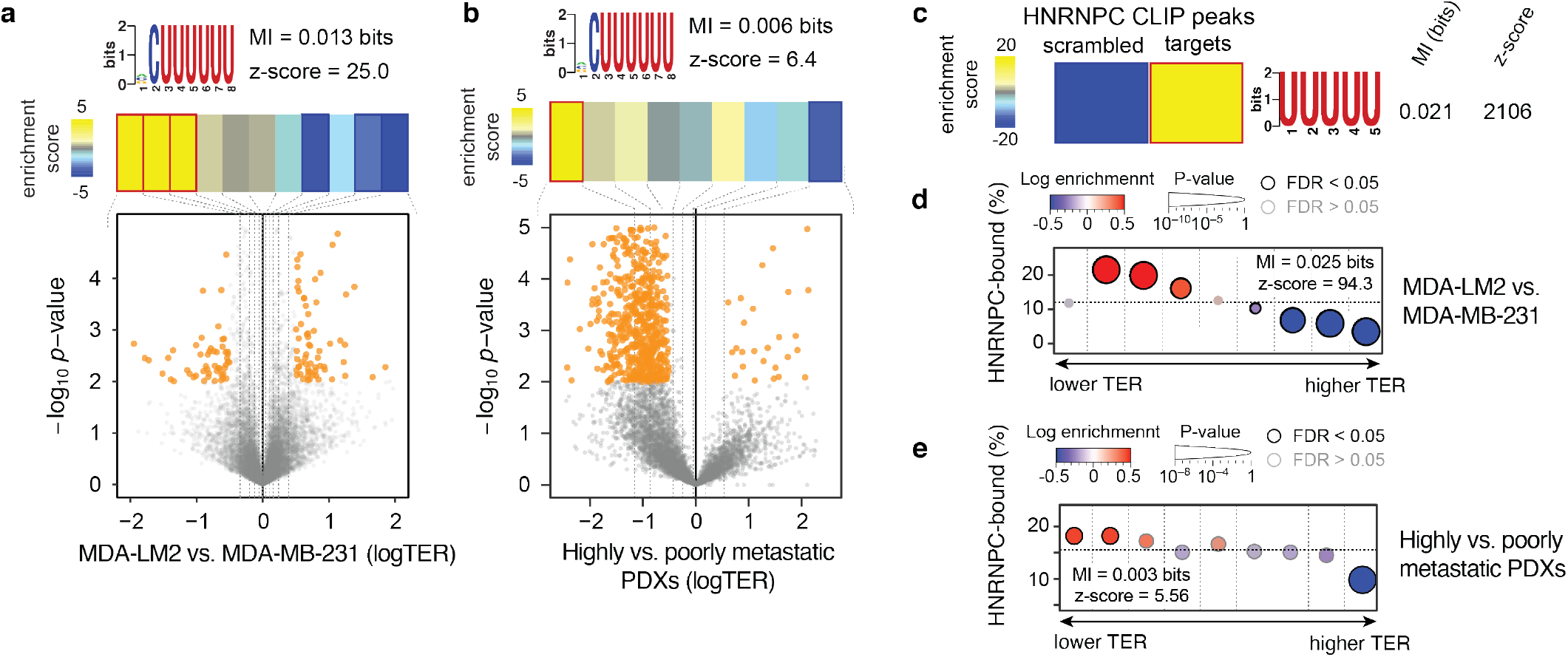
HNRNPC target mRNAs are translationally repressed in highly metastatic breast cancer cells and PDXs. **(a)** Bottom: Volcano plot showing the distribution of changes in translation efficiency ratio (logTER) in MDA-LM2 compared to parental MDA-MB-231 cells. Statistically significant (logistic regression, *p* < 0.01) observations are highlighted in orange. Top: Enrichment of the poly(U) motif in the mRNA 3’ UTRs as a function of logTER between MDA-LM2 and MDA-MB-231 cells. mRNAs are divided into equally populated bins based on their logTER (dotted vertical lines delineate the bins). Bins with significant enrichment (hypergeometric test, corrected *p* < 0.05; red) or depletion (blue) of poly(U) motifs are denoted with a bolded border. Also included are mutual information (MI) values and their associated *z*-scores. **(b)** Volcano plot showing the distribution of changes in translation efficiency in highly versus poorly metastatic breast cancer PDXs, as described for (a). **(c)** Heatmap showing the enrichment of poly(U) motifs among the HNRNPC binding sites (as determined by CLIP-seq) as compared to scrambled sequences (with di-nucleotide frequency held constant). The bolded border denotes a statistically significant enrichment (hypergeometric test, corrected *p* < 0.05; red). MI value and associated *z*-score are shown. **(d)** Enrichment of the HNRNPC target mRNAs as a function of logTER between MDA-LM2 and MDA-MB-231 cells. mRNAs are binned as in (a); the y-axis shows the frequency of the HNRNPC targets (3’ UTR-bound) that we identified in each bin (dotted horizontal line denotes the average HNRNPC target frequency across all transcripts). Bins with significant enrichment (logistic regression, FDR < 0.05; red) or depletion (blue) of HNRNPC targets are denoted with a black border. **(e)** Enrichment patterns of HNRNPC target mRNAs as a function of logTER between highly and poorly metastatic breast cancer PDXs, as in (d).

Given the extent of translational reprogramming observed in MDA-LM2 cells relative to their poorly metastatic parental line, we sought to systematically identify *cis*-regulatory elements in RNA that are significantly associated with the observed changes in translation efficiency. For this analysis, we used FIRE^19^, a mutual information-based algorithm we previously developed, to search 3’ UTRs of mRNAs for RNA motifs that are enriched in mRNAs with differential TE. FIRE identified poly(U) sequence motifs that were enriched in the 3’ UTRs of mRNAs that are translationally repressed in MDA-LM2 compared to parental cells (Fig. 1a). In order to extend our findings to other clinically relevant models of breast cancer metastasis we also performed ribosome profiling on two sets of poorly and highly metastatic human breast cancer patient-derived xenografts (PDXs)^20,21^ (Fig. S1e-f). We then compared TEs in the two highly metastatic PDXs (HCI-001 and HCI-010) to those in the two poorly metastatic PDXs (HCI-002 and STG139). We observed broad differences in the translational landscape of these PDXs, and similar to the results from the breast cancer cell lines, we observed significantly reduced translation of mRNAs with poly(U) motifs in their 3’ UTRs (Fig. 1b).

### HNRNPC controls the translation of its 3’ UTR-bound regulon

Poly(U) motifs are recognized by many RNA-binding proteins, and therefore function in a context-dependent manner^22^. To identify the most likely *trans*-factors interacting with the poly(U) sequences in translationally repressed mRNAs, we used information from the sequence context in which the poly(U) motifs are embedded. For this analysis we used DeepBind^23^, which relies on pre-trained convolutional neural networks to score target sequences of interest against a large set of RNA-binding proteins. Using this approach, we identified heterogeneous nuclear ribonucleoprotein C (HNRNPC) as the candidate most likely to bind the poly(U) motifs of interest (Fig. S1g). In agreement with a potential role for HNRNPC involvement in a translational deregulation program in metastatic breast cancer, HNRNPC was modestly but significantly downregulated in highly metastatic cells, both at the mRNA (log fold-change -0.5, *p* = 0.05, determined by RNA-seq) and protein level (log fold-change of -0.24, *p* = 0.04, determined by mass spectrometry^16^). HNRNPC ranks in the top 10% of proteins in MDA-MB-231 cells that can be detected by mass-spectrometry^24^, and therefore a slight relative decrease in protein levels corresponds to a large decrease in absolute HNRNPC abundance.

To explore the possibility that HNRNPC is a *trans*-factor that binds the identified translational regulatory poly(U) elements, we performed HNRNPC CLIP-seq^25^ in MDA-MB-231 cells. In agreement with the existing data^26^ and DeepBind predictions, poly(U) motifs were significantly enriched within HNRNPC-bound sequences (Fig. 1c). Furthermore, we detected a substantial amount of HNRNPC binding to poly(U) elements in 3’ UTRs across our own as well as previously published HNRNPC CLIP-seq datasets^26^ (Fig. S1h). Finally, if the poly(U) motifs in the 3’ UTRs of translationally repressed mRNAs are bound by HNRNPC, mRNAs that are bound by HNRNPC in their 3’ UTRs should also be translationally repressed in highly metastatic cells. To test this, we performed a gene-set enrichment analysis, using the set of HNRNPC-bound 3’ UTRs to assess their patterns of enrichment and depletion across the translational efficiency values from both breast cancer cell lines and PDXs. As shown in Fig. 1d-e (and Fig. S1i), we observed a consistent enrichment of this HNRNPC regulon among the genes with lower translational efficiency in the highly metastatic cells.

Our results above revealed a clear association between HNRNPC 3’ UTR binding and altered translational efficiency. To confirm the causal role of HNRNPC in controlling the translation of this regulon, we used CRISPR interference^27^ (CRISPRi) to knock down HNRNPC in MDA-MB-231 cells (fold-change of 0.44, determined by western blot), and used ribosome profiling to compare TEs in control and HNRNPC-deficient cells. HNRNPC knockdown affected the translational landscape in MDA-MB-231 cells, and, specifically, caused translational repression of HNRNPC target mRNAs (Fig. 2a, Fig. S2a). We found that for the most part the same mRNAs were translationally repressed in MDA-LM2 and HNRNPC knockdown cells, further highlighting the role of HNRNPC as a regulator of translational efficiency (Fig. S2b). However, HNRNPC is a predominantly nuclear protein and a known regulator of alternative splicing^28^, a fact that is reflected in our CLIP-seq data as well based on its pervasive binding to intronic sequences (Fig. S1h). Therefore, it was unclear how HNRNPC, as a nuclear protein, could impact the translation of its targets in the cytoplasm. We thus hypothesized that HNRNPC might act indirectly to control the translation of its targets.

**Figure 2.**
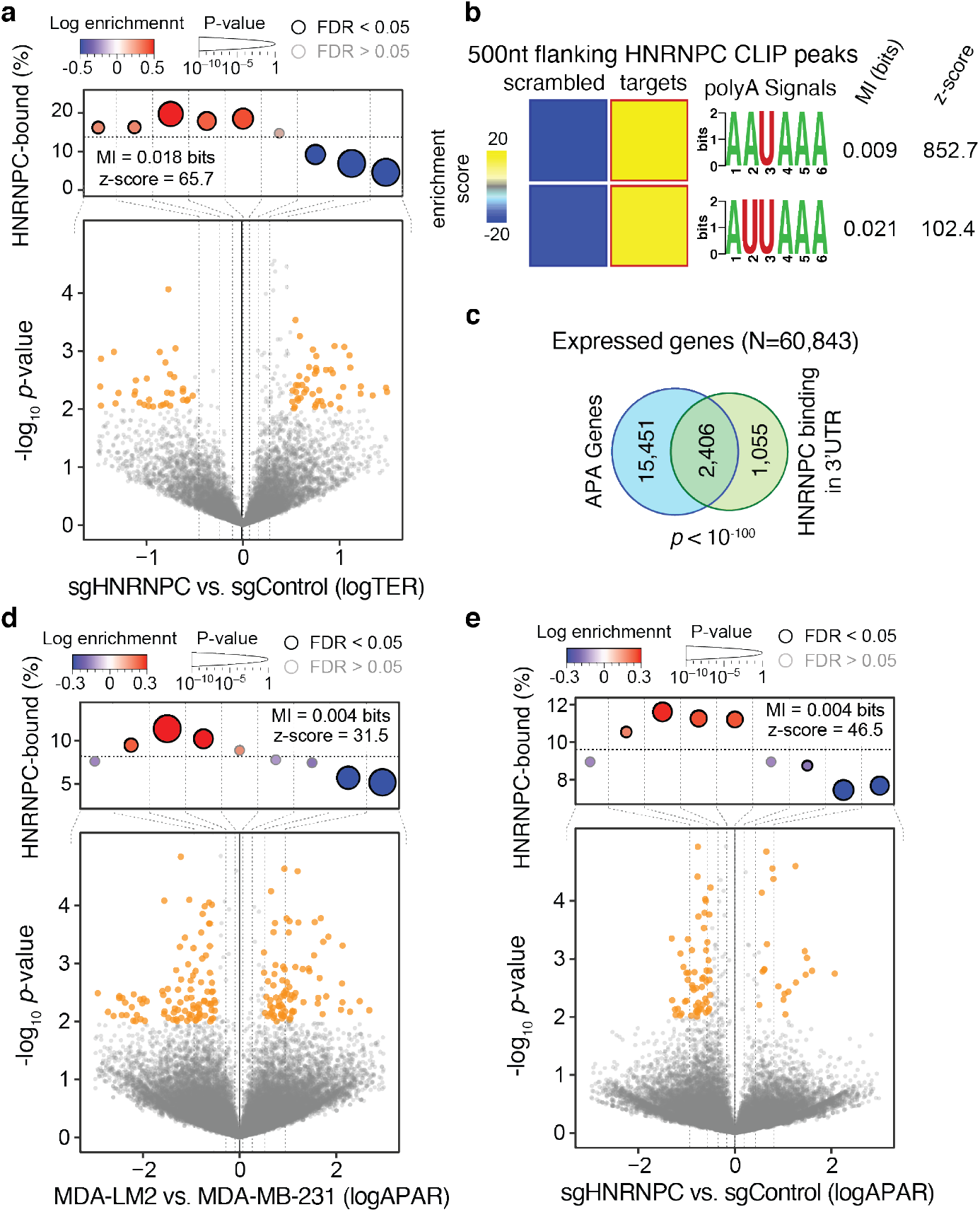
HNRNPC binding impacts the translation and alternative polyadenylation of its targets. **(a)** Bottom: Volcano plot showing the distribution of changes in translation efficiency ratio (logTER) in sgHNRNPC compared to sgControl MDA-MB-231 cells. Statistically significant (logistic regression, *p* < 0.01) observations are highlighted in orange. Top: Enrichment of the HNRNPC targets as a function of logTER between sgHNRNPC and sgControl cells. mRNAs are divided into equally populated bins according to logTER (dotted vertical lines delineate the bins); the y-axis shows the frequency of the HNRNPC targets that we identified in each bin (dotted horizontal line denotes the average HNRNPC target frequency across all transcripts). Bins with significant enrichment (logistic regression, FDR < 0.05; red) or depletion (blue) of HNRNPC targets are denoted with a black border. Also included are mutual information (MI) values and their associated *z*-scores. **(b)** Heatmaps showing the enrichment of canonical poly(A) signals in the vicinity (500 nt flanking) of HNRNPC binding peaks in 3’ UTRs (as determined by CLIP-seq). The bolded border denotes a statistically significant enrichment (hypergeometric test, corrected *p* < 0.05; red). MI values and associated *z*-scores are shown. **(c)** Venn diagram showing the overlap between HNRNPC 3’ UTR target mRNAs and mRNAs showing alternative polyadenylation. *P* value calculated using hypergeometric test. **(d)** Bottom: Volcano plot showing distribution of changes in alternative polyadenylation ratio (logAPAR, see Methods for detailed description) in MDA-LM2 compared to MDA-MB-231 cells. Top: Enrichment of the HNRNPC-bound 3’ UTRs as a function of APAR between MDA-LM2 and parental MDA-MB-231 cells; statistics as in (a). **(e)** Bottom: Volcano plot showing distribution of changes in APAR in sgHNRNPC compared to sgControl cells. Top: Enrichment of the HNRNPC-bound 3’ UTRs as a function of APAR between sgHNRNPC and sgControl cells; statistics as in (a).

### HNRNPC controls the alternative polyadenylation of its targets

To better capture the regulatory context within which HNRNPC functions, we performed a systematic search for additional *cis*-regulatory elements in the vicinity of HNRNPC binding sites on 3’ UTRs. Interestingly, as shown in Fig. 2b, we observed a highly significant enrichment of canonical poly(A) signals (AAUAAA and AUUAAA)^15^ within the 500 nucleotide flanking regions of HNRNPC CLIP-seq peaks in 3’ UTRs. This observation led us to the hypothesis that HNRNPC controls translation by regulating 3’ UTR length via alternative polyadenylation (APA) site selection. Consistently, the majority of HNRNPC 3’ UTR targets carry annotated alternative polyadenylation sites, significantly more than expected by chance (Fig. 2c). In line with this, HNRNPC has been previously implicated in the control of alternative polyadenylation in other studies, however, the mechanism through which HNRNPC impacts polyadenylation remained uncertain^29,30^. To confirm that the changes in translation efficiency we observed in highly metastatic cells coincide with alteration in poly(A) site selection, we performed mRNA 3’ end sequencing in the parental MDA-MB-231 and MDA-LM2 cells, as well as in control and HNRNPC knockdown cells. In order to measure changes in poly(A) site selection, we tabulated the number of reads that map to each annotated poly(A) site across the transcriptome. We then used a quantity we call logAPAR (log fold-change in proximal-to-distal alternative polyadenylation ratio) to identify poly(A) site switches between conditions.

To assess the statistical significance of the observed changes in alternative polyadenylation (i.e., non-zero logAPAR), we developed a novel method named APAlog. APAlog runs multinomial logistic regression to test differential usage of two or multiple poly(A) sites per transcript, and simultaneously calculates logAPAR and its associated *p*-value. It functions in three modes: (*i*) identifying transcripts with the highest overall variability in poly(A) site usage across conditions, (*ii*) comparing all non-canonical poly(A) sites to one canonical (reference) poly(A) site per transcript, and (*iii*) comparing all pairs of poly(A) sites per transcript (see Methods for details). Using APAlog, we found that HNRNPC target mRNAs undergo 3’ UTR lengthening (i.e. proximal-to-distal poly(A) site switch) in MDA-LM2 (Fig. 2d, Fig. S2c) and HNRNPC knockdown (Fig. 2e, Fig. S2d-e) cells. This observation demonstrates that HNRNPC acts as a direct mediator of an alternative poly(A) site selection program in metastatic breast cancer with broad consequences on the translational landscape.

### HNRNPC acts together with PABPC4 to control alternative polyadenylation

To obtain insights into how HNRNPC differentially controls polyadenylation of its target RNAs in parental MDA-MB-231 and MDA-LM2 cells, we immunoprecipitated HNRNPC in both cell lines and identified interacting proteins by mass spectrometry. We specifically searched for ways in which the HNRNPC interactome switches between poorly and highly metastatic cells. First, in agreement with the canonical role of HNRNPC as a splicing regulator, we detected numerous splicing factors among HNRNPC interactors (Table S1). However, when comparing the HNRNPC interactomes between poorly and highly metastatic cells, we did not observe broad changes in the interaction between HNRNPC and other splicing factors. In contrast, we found that a group of proteins implicated in mRNA transport from the nucleus was significantly depleted from the HNRNPC interactome in MDA-LM2 cells (Fig. S3a). Upon closer inspection of the proteins in this set, we noted multiple poly(A)-binding proteins that play canonical roles in the mRNA nuclear export cascade^31^. We found that PABPC4 and PABPN1 were among the depleted HNRNPC interactors in MDA-LM2 cells (Fig. 3a, Fig. S3b). We individually confirmed an RNA-dependent interaction between HNRNPC and PABPC4 or PABPN1 in MDA-MB-231 cells by co-immunoprecipitation and western blotting (Fig. 3b). Next, in order to assess which, if any, of these factors acts in concert with HNRNPC to regulate poly(A) site selection, we depleted PABPC4 and PABPN1 in MDA-MB-231 cells using CRISPRi and employed 3’ end RNA-seq to compare the APA landscapes in control and knockdown cells. We found that PABPC4 knockdown, but not PABPN1 knockdown, resulted in APA changes similar to those observed in HNRNPC-deficient cells (Fig. S3c-d). Importantly, PABPC4 was also downregulated in MDA-LM2 cells compared to parental MDA-MB-231 cells at both the mRNA (log fold-change -0.9, *p* = 0.02, determined by RNA-seq) and protein level (log fold-change -0.5, *p* < 0.002, determined by mass scpetrometry^16^).

**Figure 3.**
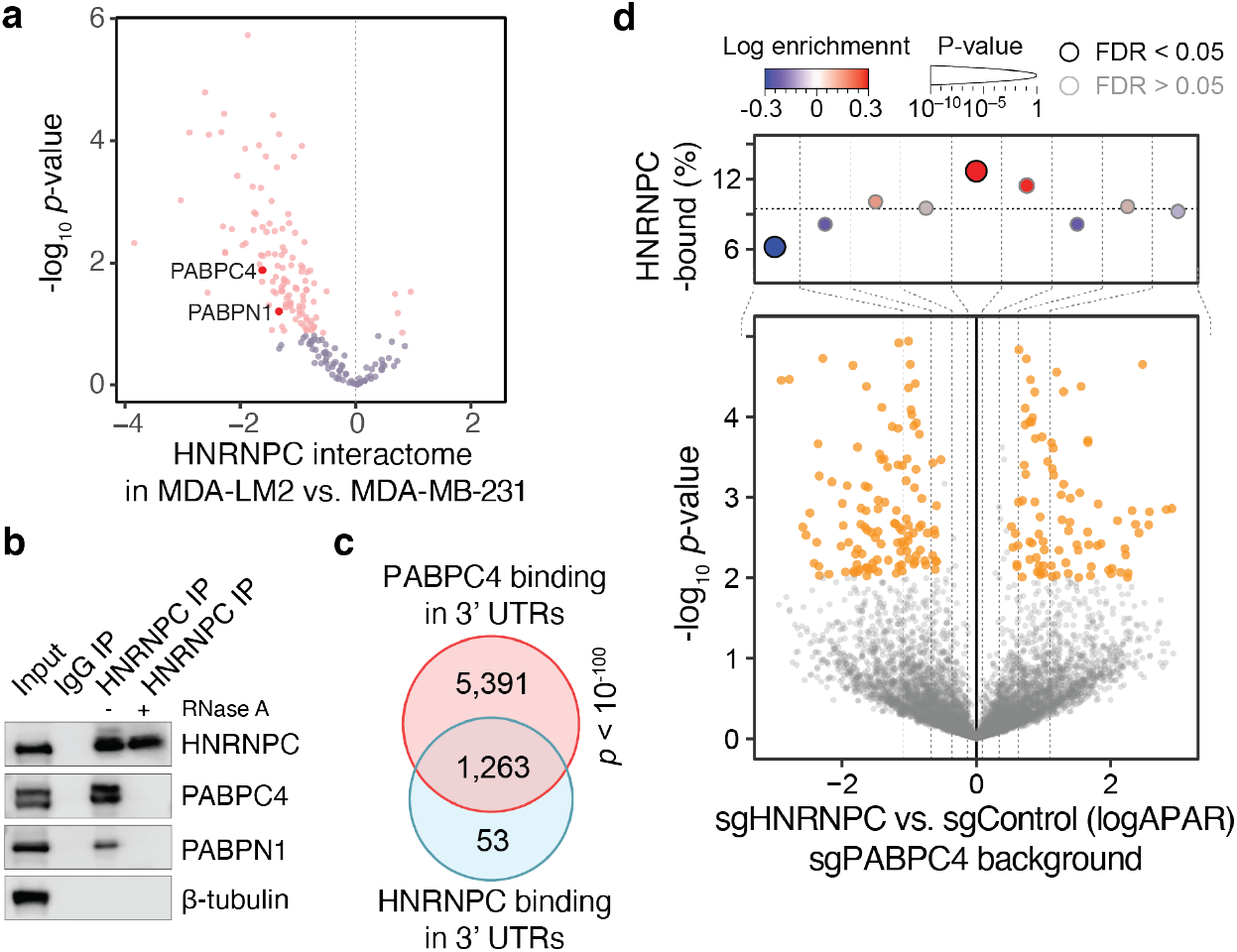
PABPC4 acts in concert with HNRNPC to control alternative polyadenylation of its target mRNAs. **(a)** Volcano plot showing the distribution of changes in relative protein interaction with HNRNPC (as determined by HNRNPC or control (isotype IgG) co-immunoprecipitation and mass spectrometry) in MDA-LM2 compared to MDA-MB-231 cells. Statistically significant (FDR-adjusted *p* value < 0.25) observations are highlighted in pink. **(b)** Co-immunoprecipitations of HNRNPC or control IgG were analyzed by western blotting. RNase A was included in the lysates where indicated. **(c)** Venn diagram showing the overlap between HNRNPC and PABPC4 3’ UTR targets (as determined by CLIP-seq and PAPER-CLIP, respectively). *P* value calculated using hypergeometric test. **(d)** Bottom: Volcano plot showing distribution of changes in alternative polyadenylation ratio (logAPAR) in HNRNPC/PABPC4 double knockdown compared to PABPC4 knockdown MDA-MB-231 cells. Top: Enrichment of the HNRNPC-bound 3’ UTRs as a function of APAR between sgHNRNPC/sgPABPC4 and sgControl/sgPABPC4 cells; statistics as in Fig. 2a.

We recently introduced PAPERCLIP^32^, a CLIP-based method which systematically identifies the targets of specific poly(A) binding proteins. To further investigate the HNRNPC-PABPC4 regulon in cells, we performed PABPC4 PAPERCLIP in MDA-MB-231 cells. First, as expected, and similar to our observations for HNRNPC, we noted that PABPC4 peaks were significantly enriched in the vicinity of canonical poly(A) signals (Fig. S3e). Moreover, we observed that the majority (96%) of HNRNPC 3’ UTR targets were also bound by PABPC4 *in vivo* (Fig. 3c). To confirm that HNRNPC and PABPC4 act in concert to control the APA of HNRNPC targets, we performed 3’ end RNA-seq comparing HNRNPC/PABPC4 double knockdown cells to PABPC4 knockdown alone. Unlike Fig. 2e, in this comparison, we did not observe the significant proximal-to-distal switching that we had observed in HNRNPC and PABPC4 single knockdown cells (Fig. 3d, Fig. S3f). This finding indicates that the regulatory function of HNRNPC in poly(A) site selection is contingent on PABPC4 expression, and demonstrates an epistatic interaction between these two genes.

### The Argonaute-mediated RNA interference pathway targets the HNRNPC regulon

Thus far, we have demonstrated that HNRNPC 3’ UTR target mRNAs undergo 3’ UTR lengthening in MDA-LM2 and HNRNPC-deficient cells. We reasoned that these extended 3’ UTRs carry translationally repressive *cis*-regulatory elements, such as miRNA binding sites. Consistent with this hypothesis, we observed a significant overlap between HNRNPC-bound extended 3’ UTRs and miRNA/argonaute (AGO2) targets^33^ (Fig. 4a). MicroRNAs are known repressors of mRNA stability and translation^34^. We observed, accordingly, that mRNAs both bound by HNRNPC and containing functional miRNA binding sites^33^ had significantly lower TE than non-target mRNAs in MDA-LM2 compared to parental MDA-MB-231 and in HNRNPC-deficient compared to control cells (Fig. S4a-b). Furthermore, translationally repressed mRNAs in MDA-LM2 and HNRNPC-deficient cells were enriched among AGO2 targets (Fig. 4b, Fig. S4c). To validate that miRNAs contribute to the translational repression of HNRNPC targets via an AGO2-mediated mechanism, we performed ribosome profiling in control and HNRNPC-depleted MDA-MB-231 cells, both in control and AGO2 knockdown backgrounds. We found that HNRNPC-depletion driven translational repression was contingent on AGO2 expression (Fig. S4d). Similarly, when we compared TEs in control and AGO2 knockdown cells, we found that mRNAs with higher TEs in AGO2-depleted cells were enriched in HNRNPC-bound transcripts (Fig. 4c).

**Figure 4.**
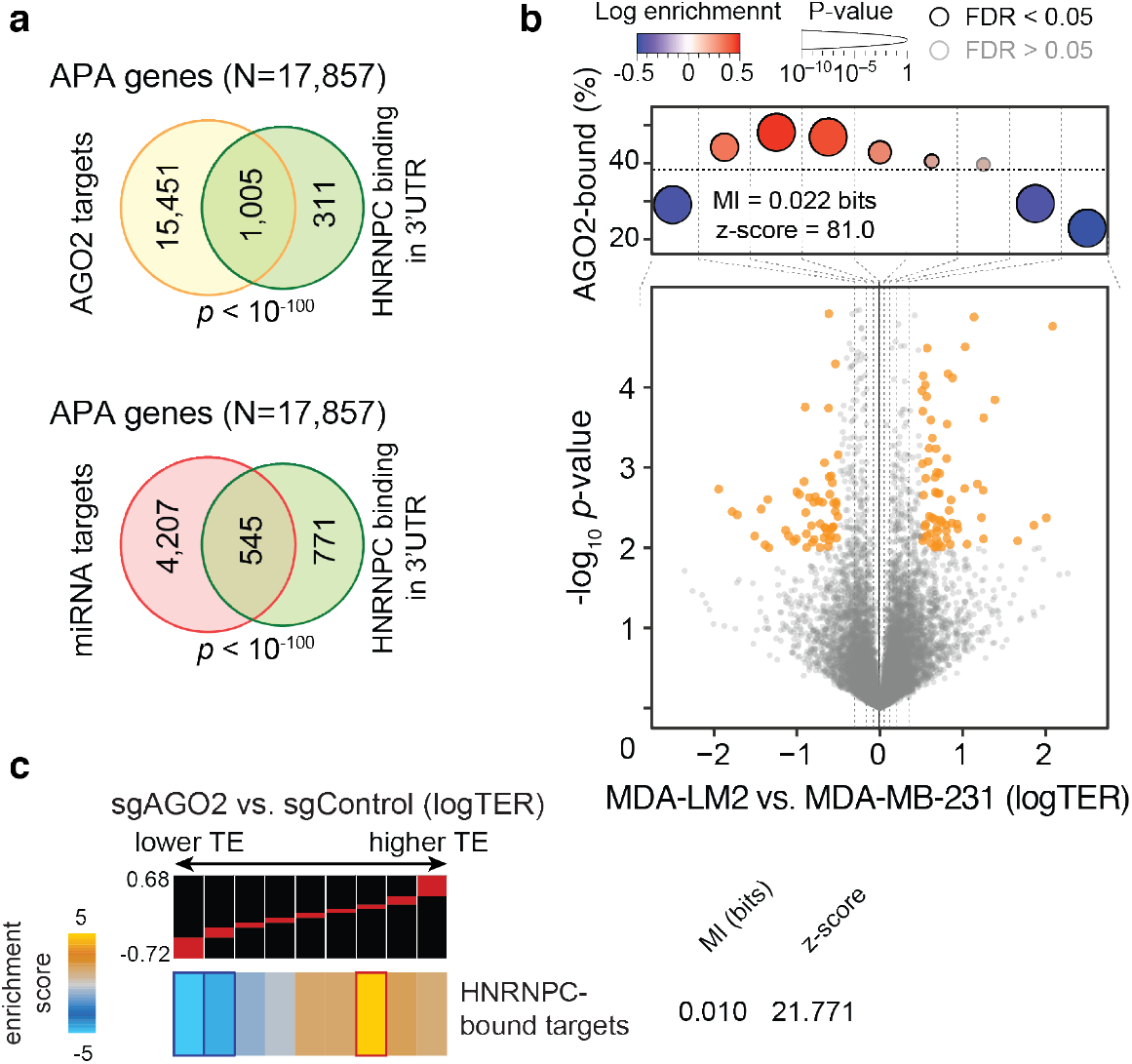
HNRNPC target mRNAs undergo miRNA-mediated translational repression. **(a)** Venn diagram showing the overlap between HNRNPC 3’ UTR targets and AGO2 bound mRNAs (top) or miRNA target mRNAs (bottom), as determined by CLIP-seq analyses. *P* values calculated using hypergeometric test. **(b)** Bottom: Volcano plot showing distribution of changes in translation efficiency ratio (logTER) in sgAGO2 compared to sgControl MDA-MB-231 cells. Top: Enrichment of the AGO2 targets as a function of logTER between sgAGO2 and sgControl cells; statistics as in Fig. 2a. **(c)** Enrichment of the HNRNPC targets as a function of logTER between sgAGO2 and sgControl cells. mRNAs are distributed into equally populated bins according to their logTER (the red bars on the black background show the range of values in each bin). Bins with significant enrichment (hypergeometric test, corrected *p* < 0.05; red) or depletion (blue) of HNRNPC targets (3’ UTR-bound) are denoted with a bolded border. Also included are mutual information (MI) value and its associated *z*-score.

### HNRNPC and PABPC4 act as suppressors of metastasis in xenograft models

We have thus far used comparisons of poorly and highly metastatic cells to uncover a previously unknown mechanism of translational control that is compromised in highly metastatic cells. However, on its own, differential activity of this pathway does not imply a functional role in metastatic progression. To assess this possibility, we used a xenograft mouse model of metastasis to measure the impact of perturbing this HNRNPC-mediated pathway on the metastatic capacity of the cell. We first performed lung colonization assays in NOD *scid* gamma mice by intravenously injecting control and HNRNPC-knockdown cells, constitutively expressing luciferase. We monitored the metastatic burden in the lungs of these mice by *in vivo* bioluminescence imaging, and observed an over ten-fold increase in lung colonization capacity induced by HNRNPC knockdown (Fig. 5a). To ensure that these findings are generalizable to other genetic backgrounds, we performed metastatic lung colonization assays with HCC1806 breast cancer cells and observed a consistent increase in the metastatic capacity of these cells upon HNRNPC down-regulation (Fig. 5b).

**Figure 5.**
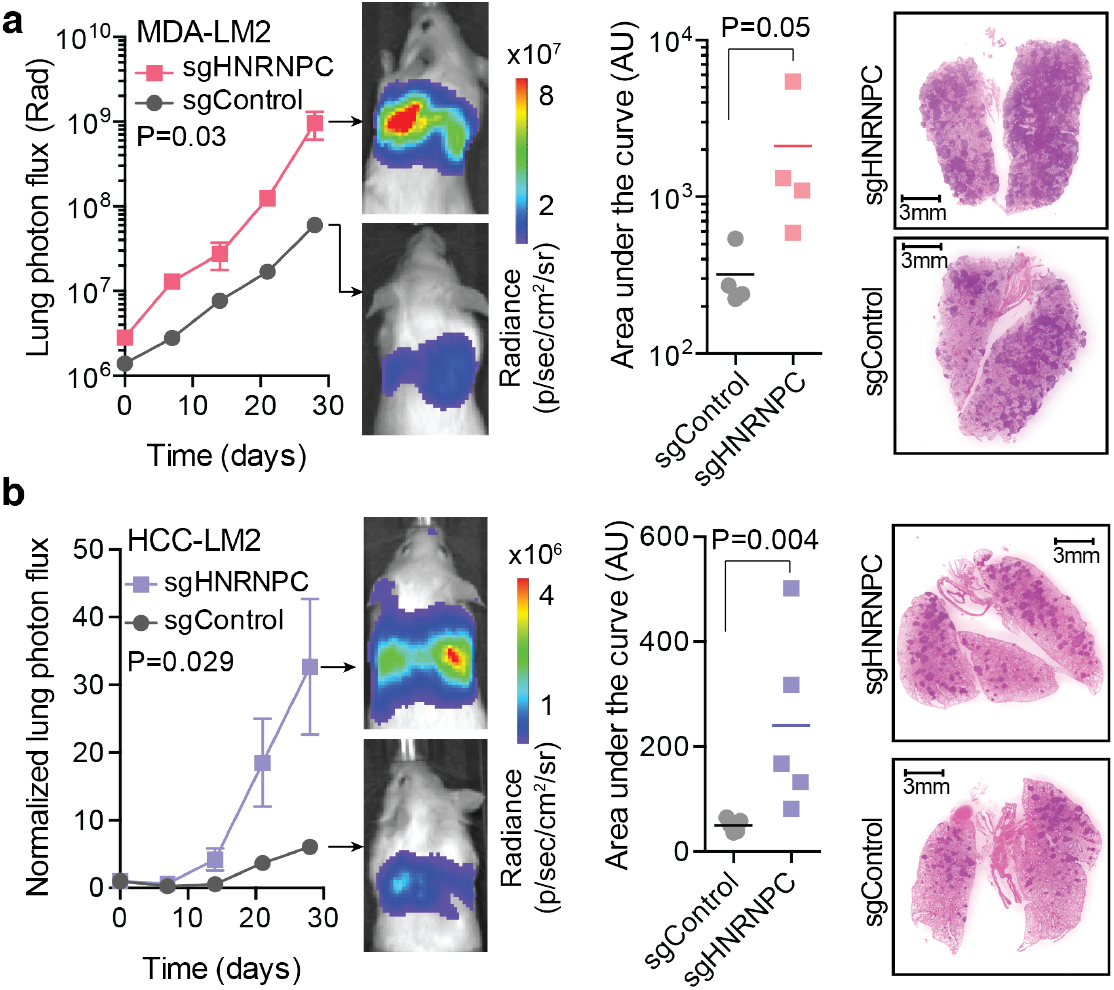
HNRNPC levels impact *in vivo* metastatic colonization of breast cancer cells. **(a)** MDA-LM2 cells stably expressing sgHNRNPC or sgControl were injected via tail vein into NSG mice. Bioluminescence was measured at the indicated times (*p* value calculated using two-way ANOVA); area under the curve was measured at the final time point (*p* value calculated using one-tailed Mann-Whitney *U*-test). Lung sections were stained with H&E (representative images shown). *n* = 4-5 mice per cohort. **(b)** HCC1806-LM2 cells stably expressing sgHNRNPC or sgControl were injected via tail vein into NSG mice. Bioluminescence was measured at the indicated times (*p* value calculated using two-way ANOVA); area under the curve was measured at the final time point (*p* value calculated using one-tailed Mann-Whitney *U*-test). Lung sections were stained with H&E (representative images shown). *n* = 4-5 mice per cohort.

In its canonical role, HNRNPC acts as a regulator of RNA splicing, and its function in metastasis may in fact be a consequence of these parallel regulatory programs. To assess this possibility, we sought to independently test PABPC4, which acts in concert with HNRNPC to control the APA of its targets. For this, we compared the metastatic lung colonization by PABPC4 knockdown, as well as PABPC4/HNRNPC double knockdown and control MDA-MB-231 cells (Fig. S5a). In line with PABPC4 controlling the APA of a metastasis-associated mRNA regulon, PABPC4-depleted cells showed significantly increased metastatic potential when compared to control cells. More importantly, knocking down HNRNPC in the PABPC4 knockdown background did not result in an increase metastatic potential of cells. This is consistent with HNRNPC and PABPC4 acting as components of the same regulatory pathway, and showing an epistatic genetic interaction.

### PDLIM5 acts downstream of HNRNPC to suppress metastasis

To better understand how the deregulation of APA and translation efficiency leads to increased metastatic potential, we sought to identify relevant targets downstream of the HNRNPC-PABPC4 regulatory axis. First, to complement our ribosome profiling results, we also compared the protein abundances in control and HNRNPC knockdown cells using TMT-MS. We observed that (*i*) consistent with its role in translational control, a large number of proteins were dysregulated upon HNRNPC depletion (Fig. S6a), and (*ii*) changes in the protein landscape of HNRNPC knockdown cells were significantly correlated with those between highly and poorly metastatic cells (Fig. S6b). In other words, a significant portion of changes in the translational efficiency of MDA-LM2 cells relative to parental MDA-MB-231 can be explained by lower HNRNPC activity. Furthermore, gene-set enrichment analysis of this data revealed that HNRNPC knockdown caused the downregulation of proteins interacting with SH3 (Scr homology 3) domain proteins and actin filaments, among other gene ontology terms (Fig. S6c).

In order to identify genes that are part of this HNRNPC regulon and act downstream of this pathway to influence metastatic progression, we systematically integrated the datasets comparing poorly and highly metastatic cells, as well as HNRNPC knockdown and control cells. We specifically searched for mRNAs that (*i*) were translationally repressed and (*ii*) demonstrated proximal-to-distal poly(A) site switching in MDA-LM2 and HNRNPC-deficient cells. We focused on transcripts that were bound by HNRNPC and PABPC4 to identify the direct downstream targets (Fig. 6a). This approach nominated PDLIM5, a member of cytoskeleton-associated protein family^35^, as a robust target of this HNRNPC-mediated pathway.

**Figure 6.**
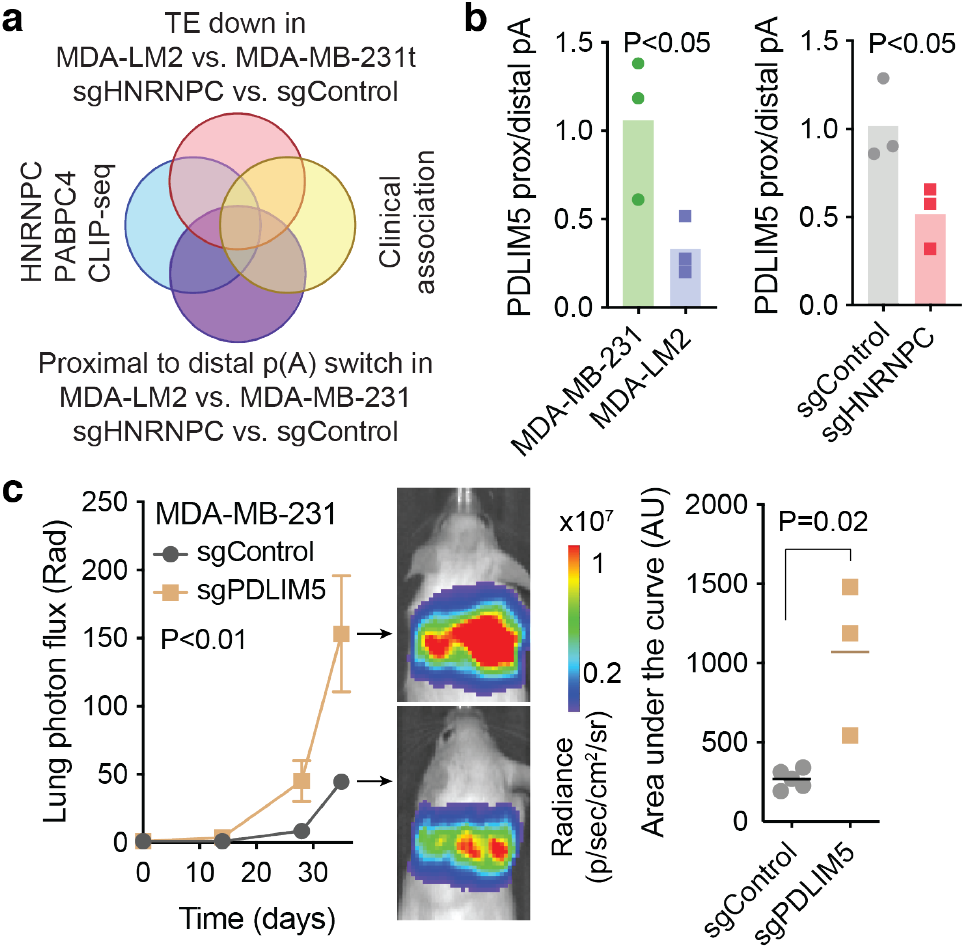
PDLIM5 acts downstream of HNRNPC to suppress breast cancer metastasis. **(a)** Venn diagram illustrating the overlap of the selected datasets for target prioritization. **(b)** Quantification of relative PDLIM5 proximal to distal poly(A) site usage in MDA-MB-231 and MDA-LM2 cells (left) or sgControl and sgHNRNPC cells (right), as determined by isoform-specific RTqPCR. *P* values calculated using one-tailed Mann-Whitney *U*-test. **(c)** MDA-MB-231 cells stably expressing sgPDLIM5 or sgControl were injected via tail vein into NSG mice. Bioluminescence was measured at the indicated times (*p* value calculated using two-way ANOVA); area under the curve was measured at the final time point (*p* value calculated using one-tailed Mann-Whitney *U*-test). *n =* 3 and 5 mice per cohort.

In agreement with our ribosome profiling and TMT-MS data (Fig. S6d), we detected lower levels of PDLIM5 protein in MDA-LM2 and HNRNPC-deficient cells, as compared to control MDA-MB-231 cells (Fig. S6e-f). In contrast, *PDLIM5* mRNA abundance was similar in these conditions, which is consistent with PDLIM5 protein levels being regulated at the translational level (Fig. S6g). We also confirmed by isoform specific RT-qPCR the proximal-to-distal poly(A) site switch for *PDLIM5* mRNA (Fig. 6b). To assess the role of PDLIM5 in breast cancer progression, we performed *in vivo* lung colonization assays with control and PDLIM5-deficient MDA-MB-231 cells. As shown in Fig. 6c, PDLIM5 knockdown led to a significant increase in metastatic lung colonization of xenografted mice.

### The HNRNPC-PABPC4 regulatory axis is associated with clinical outcomes in breast cancer patients

To confirm that our findings in xenograft models are generalizable to human disease, we performed clinical association studies in publicly available datasets from breast cancer patients. We found that lower HNRNPC expression in breast cancer tumors was significantly associated with lower overall, disease-free, and distant metastasis-free survival, both in individual cohorts and in meta-analyses (Fig. 7a-c, Fig. S7a-c). We detected that HNRNPC expression was negatively associated with tumor stage (Fig. 7d), and the presence of metastasis (Fig. 7e), but not with the tumor subtype (Fig. S7d). HNRNPC expression remained a significant covariate in a Cox proportional hazards model even after controlling for other known prognostic metrics, such as tumor stage or received treatment (Fig. S7e).

**Figure 7.**
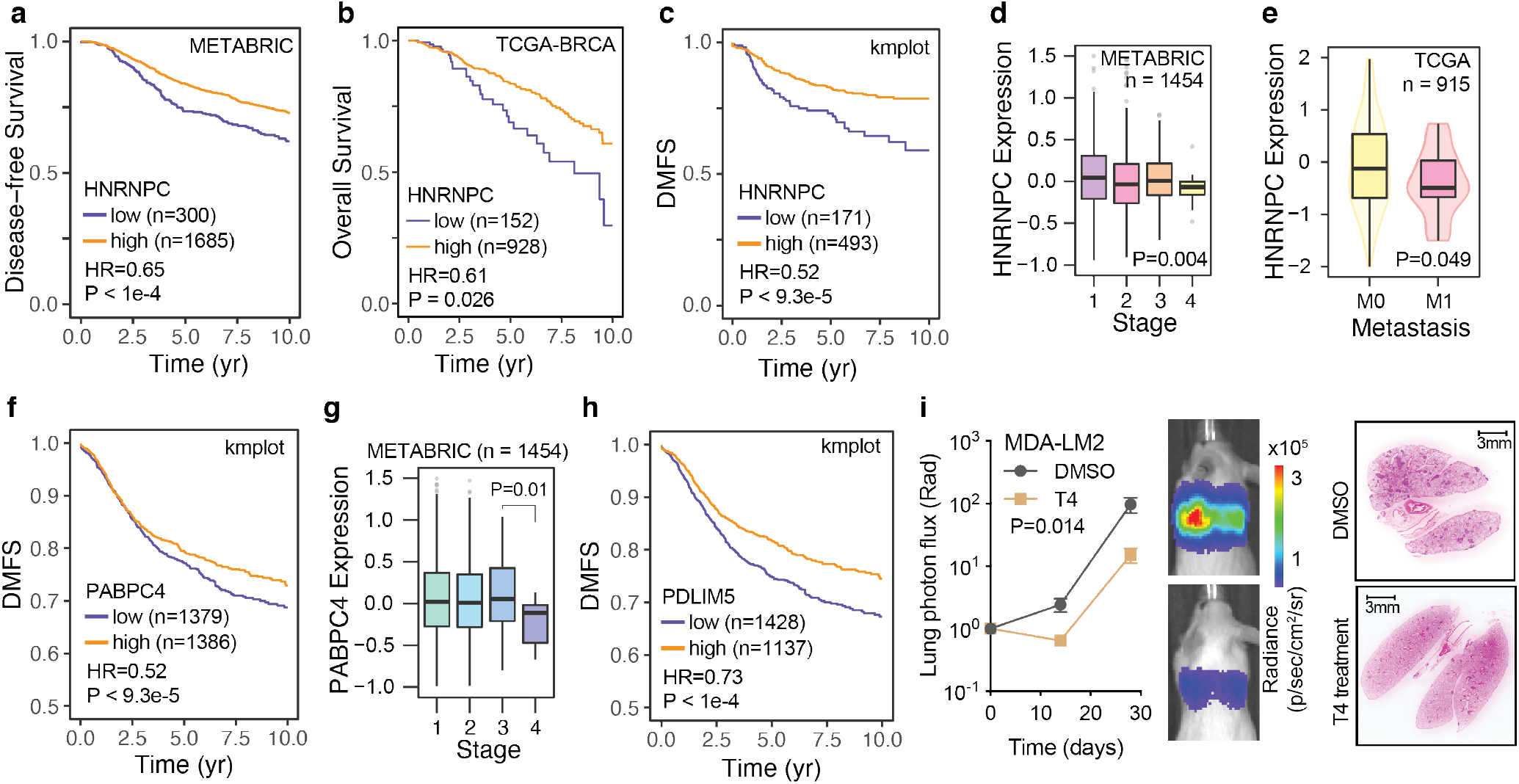
HNRNPC expression is associated with clinical outcomes in breast cancer patients. **(a)** Kaplan-Meier survival curve showing association between tumor HNRNPC levels and disease-free survival in the METABRIC cohort. **(b)** Kaplan-Meier survival curve showing association between tumor HNRNPC levels and overall survival in the TCGA-BRCA cohort. **(c)** Kaplan-Meier survival curve showing association between tumor HNRNPC levels and distant metastasis-free survival (DMFS) in a collection of breast cancer patient cohorts. **(d)** HNRNPC mRNA levels across breast cancer tissue stages I-IV in the METABRIC cohort. *P* value calculated using ANOVA. **(e)** HNRNPC mRNA levels in non-metastatic (M0) and metastatic (M1) breast tumors in the TCGA-BRCA cohort. *P* value calculated using Mann-Whitney *U*-test. **(f)** Kaplan-Meier survival curve showing association between tumor PABPC4 levels and distant metastasis-free survival (DMFS) in a collection of breast cancer patient cohorts. **(g)** PABPC4 mRNA levels across breast cancer tissue stages I-IV in the METABRIC cohort. *P* value calculated using ANOVA. **(h)** Kaplan-Meier survival curve showing association between tumor PDLIM5 levels and distant metastasis-free survival (DMFS) in a collection of breast cancer patient cohorts. Hazard ratios (HR) and *p* values (calculated using log-rank test) are shown. **(i)** MDA-LM2 cells treated with T4 or vehicle control (DMSO) at 3 μM for 6 hours were injected via tail vein into NSG mice. Bioluminescence was measured at the indicated times (*p* value calculated using two-way ANOVA). Lung sections were stained with H&E (representative images shown). *n =* 4-5 mice per cohort.

As we found HNRNPC to act upstream of a metastasis-suppressive translational program, we identified a set of HNRNPC mRNA targets, translationally repressed and undergoing proximal-to-distal poly(A) site switching in MDA-LM2 cells to define a translational HNRNPC target signature. We observed that in proteomic datasets from breast cancer patients (CPTAC), lower protein levels of the HNRNPC signature, as an aggregate, were significantly associated with lower overall and progression-free survival (Fig. S7f-g). In line with PABPC4 acting together with HNRNPC, lower PABPC4 expression was associated with worse prognostic metrics and disease progression in breast cancer patient cohorts (Fig. 7f-g, Fig. S7h). Furthermore, in agreement with PDLIM5 being a functional effector downstream of the HNRNPC-PABPC4 axis, PDLIM5 expression was also associated with breast cancer patient survival (Fig. 7h, Fig. S7i-j).

Finally, we asked whether the impact of the HNRNPC-PABPC4 deficiency in highly metastatic cells could be reversed as a potential therapeutic strategy to prevent metastasis. Recently, a target agnostic chemical screen was used to identify small molecules that impact alternative polyadenylation or transcription termination^36^. We chose to test T4, a drug that was reported to induce distal-to-proximal poly(A) site switch^36^, i.e. the opposite of the observed 3’ UTR lengthening of HNRNPC targets in MDA-LM2 cells. We first confirmed that treating MDA-MB-231 cells with 5 μM T4 for 6 hours induced predominantly distal-to-proximal poly(A) site switching, as assessed by 3’ end RNA-seq (Fig. S7k). Importantly, we observed that T4 treatment can reverse the impact of HNRNPC knockdown on APA site selection (Fig. S7l-m).

These results suggested that T4 could potentially counteract the 3’ UTR lengthening of HNRNPC targets, and compromise the pro-metastatic program instigated by HNRNPC deficiency. To test this possibility, we first performed dose-response measurements of T4 in MDA-LM2 cells (Fig. S5n), and treated these cells with the IC20 concentration of the drug (3 μM) or vehicle control for 6 hours. We then performed lung colonization assays to measure changes in the metastatic capacity of cells treated with T4. As shown in Fig. 7i, T4 treated MDA-LM2 cells showed significantly reduced metastatic capacity as compared to vehicle treated cells. This observation indicates that reversing the regulatory consequences of HNRNPC down-regulation can restore the metastasis-suppressive activity of its target regulon.

## DISCUSSION

In this study, we show that increased metastatic potential in breast cancer cell lines and PDXs is accompanied by a broad remodeling of the translational landscape, as demonstrated by genome-wide TE measurements derived from ribosome profiling. Using unbiased computational approaches, we discovered that translationally downregulated mRNAs in highly metastatic breast cancer cell lines and PDXs showed enriched poly(U) motifs in their 3’ UTRs. U-rich sequences are recognized by a number of RNA binding proteins (RBPs), including ELAVL1, TIA1, TIAL1, U2AF2, CPEB2, CPEB4, and HNRNPC^22^. Interactions between poly(U) motifs and these RBPs is context dependent, and has been linked to alternative splicing and polyadenylation, altered mRNA stability and, to a lesser extent, translation^15^. Focusing on the poly(U) stretches in the 3’ UTRs of translationally repressed mRNAs, we discovered a link between HNRNPC binding to these sequence elements and translational control of the bound transcript. We showed that this mechanism relies on control of the alternative polyadenylation of HNRNPC target mRNAs.

We found that HNRNPC deficiency resulted in increased usage of distal poly(A) sites of target transcripts, leading to 3’ UTR lengthening of HNRNPC targets, consistent with previous reports^29,30^. It was suggested that HNRNPC masks strong distal poly(A) sites, thereby promoting usage of weaker proximal sites^29^. In line with this, we found that canonical poly(A) signals were in close proximity to HNRNPC-bound poly(U) motifs. Interplay between alternative polyadenylation and splicing has been reported^15^, supporting the role of HNRNPC in poly(A) site choice while its primary function has been associated with splicing^28^. We could counteract the proximal-to-distal poly(A) site switch caused by HNRNPC deficiency with T4, a small molecule, promoting a distal-to-proximal switch^36^. While the mechanism of action of T4 is not completely clear, it was found to alter the expression levels of multiple splicing and cleavage and polyadenylation factors^36^, emphasizing the interplay between the two pathways.

By comparing HNRNPC interactomes in highly and poorly metastatic cells, we identified a decreased interaction between HNRNPC and factors implicated in mRNA export from the nucleus, including several poly(A) binding proteins. We found that PABPC4 bound HNRNPC targets *in vivo* and showed an epistatic genetic interaction with HNRNPC in controlling alternative polyadenylation. Similar to other cytoplasmic poly(A) binding proteins, PABPC4 is a nucleus-cytoplasm shuttling factor, and is known to have context dependent functions, overlapping those of other poly(A) binding proteins^31^. PABPC4 is critical for the differentiation of erythroid cells, via an interplay between AU-rich elements in 3’ UTR of target mRNAs and the shortening of poly(A) tails^37^. Poly(A) tail shortening is a well-known mechanism in promoting mRNA decay and downregulating translation^38^. It is possible that reduced PABPC4 expression in highly metastatic cells contributes to the lower TE of joint HNRNPC-PABPC4 targets via poly(A) tail shortening. Interestingly, in colon cancer, higher PABPC4 expression was associated with better clinical outcomes^39^, consistent with our observations in breast cancer clinical data.

Our data support the hypothesis that the long form 3’ UTRs of HNRNPC target mRNAs harbor a greater number of miRNA binding sites and thus are more susceptible to translational repression via argonaute-mediated RNA interference^10,34^. Although some studies suggest that miRNAs have a limited impact on global translational repression and destabilization of APA targets^40,41^, our data highlights a case where a subset of mRNAs—HNRNPC targets with 3’ UTR lengthening in highly metastatic cells—undergo AGO2-dependent translational repression. Furthermore, HNRNPC targets inherently contain a poly(U) stretch in their 3’ UTR, and in conditions where HNRNPC levels are low, this might favor interaction with other poly(U) binding RBPs and lead to translational repression in the cytoplasm.

Recent data suggests that HNRNPC binding to RNA is affected by post-transcriptional RNA modifications, such as *N*^*6*^-methyladenosine (m6A). Dynamic RNA methylation can impact the secondary structure of modified RNA sequences, thereby altering the accessibility for RBP binding^42–44^. In MALAT1 RNA, m6A modification of a specific residue reduces base pairing with an adjacent poly(U) stretch, promoting HNRNPC binding to this poly(U) region^42^. Canonical poly(A) signals are (A)-rich motifs, and we have found that HNRNPC-bound poly(U) sequences are often in the vicinity of poly(A) signals. The sequence complementarity between poly(U) stretches and poly(A) signals makes their interaction plausible, particularly in an HNRNPC-deficient context. Therefore, it is conceivable that m6A modification of regions adjacent to HNRNPC binding sites may add another layer of regulation to the translational control pathway described here, as the dynamics between RNA secondary structure and HNRNPC binding is likely to affect alternative polyadenylation.

We found that HNRNPC deficiency results in altered PDLIM5 mRNA polyadenylation site usage and reduced translation, and that PDLIM5 knockdown phenocopied the pro-metastatic phenotype of HNRNPC-depleted cells. PDLIM5 (PDZ and LIM domain 5) is a member of cytoskeleton-associated protein family, implicated in cell-cell, cell-extracellular matrix interactions, and cell migration^35^. It also participates in the mechanosensing cascade via YAP/TAZ signalling^45^. PDLIM5 is phosphorylated by AMPK, and this modulates its function in cell migration^46^. Interestingly, these cellular and molecular functions were enriched among the downregulated proteins in HNRNPC-depleted cells. Altered interactions with the tumour microenvironment and increased migratory potential are among the major routes to metastasis, and we propose that PDLIM5 repression plays a role in the pro-metastatic program induced by HNRNPC deficiency.

We have uncovered an intricate gene regulatory program at the intersection of alternative polyadenylation and translational control mediated by HNRNPC and PABPC4 that plays a metastasis suppressing role in breast cancer. Our clinical association analyses suggest that HNRNPC expression, along with that of its regulon, could be used as a prognostic metric for disease progression. We also provide evidence that HNRNPC low tumours could benefit from therapeutic strategies targeting alternative polyadenylation.

## METHODS

### Cell culture

All cells were cultured in a 37°C 5% CO2 humidified incubator. The MDA-MB-231 (ATCC HTB-26) breast cancer cell line, its highly metastatic derivative, MDA-LM2^17^, and 293T cells (ATCC CRL-3216) were cultured in DMEM high-glucose medium supplemented with 10% FBS, glucose (4.5 g/L), L-glutamine (4 mM), sodium pyruvate (1 mM), penicillin (100 units/mL), streptomycin (100 μg/mL) and amphotericin B (1 μg/mL) (Gibco). The HCC1806-LM2 cell line (an *in vivo* selected highly lung metastatic derivative of the HCC1806 breast cancer line (ATCC CRL-2335)) was cultured in RPMI-1640 medium supplemented with 10% FBS, glucose (2 g/L), L-glutamine (2 mM), 25 mM HEPES, penicillin (100 units/mL), streptomycin (100 μg/mL) and amphotericin B (1 μg/mL) (Gibco). All cell lines were routinely screened for mycoplasma with a PCR-based assay.

### T4 treatment

T4 (Enamine EN300-7536403) was dissolved in DMSO at 10 mM stock concentration. For 3’-end RNA-seq, cells were treated with 5 μM T4 or an equivalent amount of DMSO for 6h, prior to RNA extraction. For dose-response measurements, the cells were treated with T4 at indicated concentrations for 6h, after which the media was changed and the cell viability was measured 72 hours later using CellTiter-Glo Luminescent Cell Viability Assay (Promega) in 6 biological replicates. For *in vivo* metastasis assays, MDA-LM2 cells were treated with an IC20 concentration of T4 (3 μM) or DMSO control for 6h, and then immediately harvested for tail vein injections.

### CRISPRi-mediated gene knockdown

MDA-MB-231, MDA-LM2 and HCC1806-LM2 cells expressing dCas9-KRAB fusion protein were constructed by lentiviral delivery of pMH0006 (Addgene #135448) and FACS isolation of BFP-positive cells.

The lentiviral constructs were co-transfected with pCMV-dR8.91 and pMD2.D plasmids using TransIT-Lenti (Mirus) into 293T cells, following manufacturer’s protocol. Virus was harvested 48 hours post-transfection and passed through a 0.45 μm filter. Target cells were then transduced overnight with the filtered virus in the presence of 8 μg/mL polybrene (Millipore).

Guide RNA sequences for CRISPRi-mediated gene knockdown were cloned into pCRISPRia-v2 (Addgene #84832) via BstXI-BlpI sites (see Table S2 for sgRNA sequences). For double knockdown experiments, pCRISPRia-v2 plasmid was modified to construct pCRISPRia-v2-Blast, replacing puromycin acetyltransferase by blasticidin deaminase coding sequences. After transduction with sgRNA lentivirus, MDA-MB-231, MDA-LM2 and HCC1806-LM2 CRISPRi cells were selected with 1.5 μg/mL puromycin or 20 μg/mL blasticidin (Gibco). Knockdown of target genes was assessed by RT-qPCR as described below.

### RNA isolation

Total RNA for RNA-seq and RT-qPCR was isolated using the Zymo QuickRNA isolation kit with in-column DNase treatment per the manufacturer’s protocol.

### RT-qPCR

Transcript levels were measured using quantitative RT-PCR by reverse transcribing total RNA to cDNA (Maxima H Minus RT, Thermo), then using PerfeCTa SYBR Green SuperMix (QuantaBio) per the manufacturer’s instructions. HPRT1 was used as endogenous control (see Table S2 for primer sequences).

### Western Blotting

Cell lysates were prepared by lysing cells in ice-cold RIPA buffer (25 mM Tris-HCl pH 7.6, 0.15 M NaCl, 1% IGEPAL CA-630, 1% sodium deoxycholate, 0.1% SDS) containing 1X protease inhibitors (Thermo Scientific). Lysate was cleared by centrifugation at 20,000 x g for 10 min at 4°C. Samples were denatured for 10 min at 70°C in 1X LDS loading buffer (Invitrogen) and 50 mM DTT. Proteins were separated by SDS-PAGE using 4-12% Bis-Tris NuPAGE gels, transferred to nitrocellulose (Millipore), blocked using 5% BSA, and probed using target-specific antibodies. Bound antibodies were detected using HRP-conjugated secondary antibodies and ECL substrate (Pierce) or infrared dye-conjugated secondary antibodies (Licor) according to the manufacturer’s instructions. Antibodies: beta-tubulin (Proteintech 66240-1-Ig), HNRNPC (Santa Cruz sc-32308), PABPC4 (Proteintech 14960-1-AP), PABPN1 (Proteintech 66807-1-Ig).

### Ribosome profiling

Ribosome profiling was performed as previously described^47^. Briefly, approximately 10×10^6^ cancer cells were lysed in ice cold polysome buffer (20 mM Tris pH 7.6, 150 mM NaCl, 5 mM MgCl_2_, 1 mM DTT, 100 μg/mL cycloheximide) supplemented with 1% v/v Triton X-100 and 25 U/mL Turbo DNase (Invitrogen). For PDXs, snap-frozen tumors were cryoground into powder on dry ice, and then resuspended in ice-cold lysis buffer as above. The lysates were triturated through a 27G needle and cleared for 10 min at 21,000 x g at 4°C. The RNA concentration in the lysates was determined with the Qubit RNA HS kit (Thermo). Lysate corresponding to 30 μg RNA was diluted to 200 μl in polysome buffer and digested with 1.5 μl RNaseI (Epicentre) for 45 min at room temperature. The RNaseI was then quenched by 10 μl SUPERaseIN (Thermo).

Monosomes were isolated using MicroSpin S-400 HR (Cytiva) columns, pre-equilibrated with 3 mL polysome buffer per column. 100 μl digested lysate was loaded per column (two columns were used per 200 μl sample) and centrifuged 2 min at 600 x g. The RNA from the flow through was isolated using the Zymo RNA Clean and Concentrator-25 kit. In parallel, total RNA from undigested lysates were isolated using the same kit.

Ribosome protected footprints (RPFs) were gel-purified from 15% TBE-Urea gels as 17-35 nt fragments. RPFs were then end-repaired using T4 PNK (NEB) and pre-adenylated barcoded linkers were ligated to the RPFs using T4 Rnl2(tr) K227Q (NEB). Unligated linkers were removed from the reaction by yeast 5’-deadenylase (NEB) and RecJ nuclease (NEB) treatment. RPFs ligated to barcoded linkers were pooled, and rRNA-depletion was performed using riboPOOLs (siTOOLs) as per manufacturer’s recommendations. Linker-ligated RPFs were reverse transcribed with ProtoScript II RT (NEB) and gel-purified from 15% TBE-Urea gels. cDNA was then circularized with CircLigase II (Epicentre) and used for library PCR. First, a small-scale library PCR was run supplemented with 1X SYBR Green and 1X ROX (Thermo) in a qPCR instrument. Then, a larger scale library PCR was run in a conventional PCR instrument, performing a number of cycles that resulted in ½ maximum signal intensity during qPCR. Library PCR was gel-purified from 8% TBE gels and sequenced on a SE50 run on Illumina HiSeq4000 instrument at UCSF Center for Advanced Technologies.

To process the reads, the Ribo-seq reads were first trimmed using cutadapt (v2.3) to remove the linker sequence AGATCGGAAGAGCAC. The fastx_barcode_splitter script from the Fastx toolkit was then used to split the samples based on their barcodes. Since the reads contain unique molecular identifiers (UMIs), they were collapsed to retain only unique reads. The UMIs were then removed from the beginning and end of each read (2 and 5 Ns, respectively) and appended to the name of each read. Bowtie2 (v2.3.5) was then used to remove reads that map to ribosomal RNAs and tRNAs, and the remainder of reads were then aligned to mRNAs (we used the isoform with the longest coding sequence for each gene as the representative). Subsequent to alignment, umitools (v0.3.3) was used to deduplicate reads.

### RNA-seq and 3’-end RNA-seq

RNA-seq libraries (used for calculating translation efficiencies) were prepared using SMARTer Stranded Total RNA-Seq Kit v2 - Pico Input Mammalian (Takara), with 50 ng total RNA as input. 3’-end RNA-seq libraries (used for determining poly(A) site usage) were prepared using QuantSeq 3’ mRNA-Seq Library Prep Kit REV (Lexogen), with 500 ng total RNA as input. Libraries were sequenced as SE50 runs on Illumina HiSeq4000 instrument at UCSF Center for Advanced Technologies.

To compare changes in 3’ UTR usage and poly(A) site selection, we first annotated unique 3’ ends of transcripts using Gencode annotations (v33). Salmon (v0.14.1) was then used to count the number of reads that match each of the annotated ends. The normalized abundances were then tabulated and APAlog (see below) was used to perform pairwise comparisons between proximal and distal poly(A) sites between conditions.

To assess whether HNRNPC binding was associated with the observed changes in logAPAR values, proximal sites within 500 nt of annotated HNRNPC peaks (based on CLIP-seq datasets) were annotated. The behavior of these HNRNPC-associated proximal sites was then compared to the background using a Wilcoxon rank sum test. Alternatively, logAPAR values were binned into equally populated bins, and the enrichment/depletion patterns of HNRNPC-associated proximal sites was assessed as previously described^24^.

### HNRNPC CLIP-seq

CLIP-seq for endogenous HNRNPC in MDA-MB-231 cells was performed using irCLIP^48^, with the following modifications. The cells were crosslinked with 400 mJ/cm^2^ 254 nm UV. Cells were lysed in CLIP lysis buffer (1X PBS, 0.1% SDS, 0.5% sodium deoxycholate, 0.5% IGEPAL CA-630) supplemented with 1X protease inhibitors (Thermo) and SUPERaseIN (Thermo), then treated with DNase I (Promega) for 5 minutes at 37ºC. Lysate was clarified by spinning at 21,000 x g at 4ºC for 15 min. RNA-protein complexes were then immunoprecipitated from the clarified lysate using protein G Dynabeads (Thermo) conjugated to anti-HNRNPC (Santa Cruz sc-32308) for 2 hours at 4°C. Beads were washed sequentially with high stringency buffer, high salt buffer and low salt buffer. RNA-protein complexes were then nuclease treated on-bead with RNase A (Thermo), and then ligated to the irCLIP adaptor using T4 RNA ligase (NEB) overnight at 16ºC. RNA-protein complexes were then eluted from beads, resolved on a 4-12% Bis-Tris NuPAGE gel, transferred to nitrocellulose, then imaged using an Odyssey Fc instrument (Licor). Regions of interest were excised from the membrane and the RNA was isolated by Proteinase K digestion followed by pulldown with oligo d(T) magnetic beads (Thermo). The resulting RNA was then reverse transcribed using Superscript IV RT (Invitrogen) and a barcoded RT primer, purified using MyOne C1 Dynabeads (Invitrogen), and then circularized using CircLigase II (Epicentre). Two rounds of PCR were then performed to first amplify the library using adaptor-specific primers and to add sequences compatible with Illumina sequencing instruments. The libraries were then sequenced as SE50 runs on Illumina HiSeq4000 instrument at UCSF Center for Advanced Technologies.

The CTK package (CLIP toolkit^49^) was used to annotate peaks from CLIP-seq data. Reads were first collapsed using their UMIs. UMI-tools package was then used to extract the UMI followed by quality trimming (-q 15) and linker removal using cutadapt. BWA (0.7.17) was then used to align reads to the genome (hg38). CTK scripts were then used to remove PCR duplicates, parse alignments, and call peaks using a valley-seeking algorithm (with multi-testing correction). The boundaries of the resulting peaks were combined across multiple independent CLIP experiments, and their union with a previously published HNRNPC iCLIP (E-MTAB-1371) was used to define a comprehensive HNRNPC binding across the transcriptome. To identify motifs, sequences across annotated peak were extracted; control sequences were generated by scrambling the real sequences while maintaining dinucleotide sequence content. FIRE^19^ was then used to find enriched sequence motifs.

### PABPC4 PAPERCLIP

The MDA-MB-231 cells were crosslinked with 400 mJ/cm^2^ 254 nm UV. Cells were lysed in CLIP lysis buffer (1X PBS, 0.1% SDS, 0.5% sodium deoxycholate, 0.5% IGEPAL CA-630) supplemented with 1X protease inhibitors (Thermo) and SUPERaseIN (Thermo), then treated with DNase I (Promega) for 5 minutes at 37ºC. Lysates were then split in half and separately treated with medium and low dilutions of RNaseA and RNaseI (Thermo; 1/3,000 RNaseA and 1/100 RNaseI, and 1/15,000 RNaseA and 1/500 RNaseI, respectively). Lysates were then clarified by spinning at 21,000 x g at 4ºC for 15 min. Clarified lysates were pooled and RNA-protein complexes were then immunoprecipitated using protein A/G beads (Pierce) conjugated to anti-PABPC4 (Proteintech 14960-1-AP) for 2 hours at 4°C. Beads were then washed sequentially with low salt buffer, high salt buffer and PNK buffer. Protein-bound RNAs were end-repaired on beads using T4 PNK (NEB) and 3’-end-labeled with azide-dUTP using yeast poly(A) polymerase (Jena). The protein-RNA complexes were labelled with IRDye800-DBCO conjugates (LiCor). The protein-RNA complexes were then eluted from beads, resolved on a 4-12% Bis-Tris NuPAGE gel, transferred to nitrocellulose, and imaged using an Odyssey Fc instrument (LiCor). Regions of interest were excised from the membrane and the RNA was isolated by Proteinase K digestion and phenol/chloroform extraction. Eluted RNA was used for library preparation using SMARTer smRNA-Seq Kit (Takara), with following modifications. The poly(A) tailing step was omitted, and reverse transcription was performed with a custom RT primer (see Table S2). The library PCR was performed with index forward (i5) primers and universal reverse (P7) primer (see Table S2). The libraries were purified using Zymo Select-a-Size beads and sequenced as a SE50 run on Illumina HiSeq4000 instrument at UCSF Center for Advanced Technologies. The data was analyzed as in CLIP-seq.

### Tandem mass tag labelling and mass spectrometry (TMT-MS)

The cell lysates were prepared, digested and labelled using TMT10plex Isobaric Mass Tagging Kit (Thermo), as per manufacturer’s instructions. The labelling reactions were cleaned up and fractionated using Pierce High pH Reversed-Phase Peptide Fractionation Kit (Thermo).

Peptides were analyzed on a Thermo Fisher Orbitrap Fusion Lumos Tribid mass spectrometry system equipped with an Easy nLC 1200 ultrahigh-pressure liquid chromatography system interfaced via a Nanospray Flex nanoelectrospray source. Samples were injected on a C18 reverse phase column (25 cm x 75 μm packed with ReprosilPur C18 AQ 1.9 um particles). Peptides were separated by an gradient from 5 to 32% ACN in 0.02% heptafluorobutyric acid over 120 min at a flow rate of 300 nl/min. Spectra were continuously acquired in a data-dependent manner throughout the gradient, acquiring a full scan in the Orbitrap (at 120,000 resolution with an AGC target of 400,000 and a maximum injection time of 50 ms) followed by 10 MS/MS scans on the most abundant ions in 3 s in the dual linear ion trap (turbo scan type with an intensity threshold of 5000, CID collision energy of 35%, AGC target of 10,000, maximum injection time of 30 ms, and isolation width of 0.7 m/z). Singly and unassigned charge states were rejected. Dynamic exclusion was enabled with a repeat count of 1, an exclusion duration of 20 s, and an exclusion mass width of ±10 ppm. Data was collected using the MS3 method^50^ for obtaining TMT tag ratios with MS3 scans collected in the orbitrap at a resolution of 60,000, HCD collision energy of 65% and a scan range of 100-500.

Protein identification and quantification were done with Integrated Proteomics Pipeline (IP2, Integrated Proteomics Applications) using ProLuCID/Sequest, DTASelect2 and Census^51,52^. Tandem mass spectra were extracted into ms1, ms2 and ms3 files from raw files using RawExtractor^53^ and were searched against the Uniprot human protein database plus sequences of common contaminants, concatenated to a decoy database in which the sequence for each entry in the original database was reversed^54^. Search space included all fully tryptic peptide candidates with no missed cleavage restrictions. Carbamidomethylation (+57.02146) of cysteine was considered a static modification; TMT tag masses, as given in the TMT kit product sheet, were also considered static modifications. We required 1 peptide per protein and both tryptic termini for each peptide identification. The ProLuCID search results were assembled and filtered using the DTASelect program with a peptide false discovery rate (FDR) of 0.001 for single peptides and a peptide FDR of 0.005 for additional peptides for the same protein. Under such filtering conditions, the estimated false discovery rate was between zero and 0.06 for the datasets used. Quantitative analysis on MS3-based MultiNotch TMT data was analyzed with Census 2 in IP2 platform^50,55,56^. As TMT reagents are not 100% pure, we referred to the Thermo Fisher Scientific TMT product data sheet to obtain purity values for each tag and normalized reporter ion intensities. While identification reports best hit for each peptide, Census extracted all PSMs that can be harnessed to increase accuracy from reporter ion intensity variance. Extracted reporter ions were further normalized by using total intensity in each channel to correct sample amount error.

### Co-immunoprecipitation and mass spectrometry (CoIP-MS)

MDA-MB-231 and MDA-LM2 cells (10×10^6^ per replicate) were washed with ice-cold 1X PBS and lysed in nuclei lysis buffer (100 mM Tris HCl pH 7.5, 0.5% SDS, 1 mM EDTA) containing 1X protease inhibitors (Thermo Scientific) on ice for 10 min. The lysates were then diluted with 4 volumes of IP dilution buffer (62.5 mM Tris HCl pH 7.5, 187.5 mM NaCl, 0.625% Triton X-100, 1 mM EDTA) with protease inhibitors and passed through a 25G needle several times. The lysates were cleared 10 min at 21,000 g at +4°C and used for IP.

For Co-IP/MS analysis, HNRNPC antibody was covalently bound to the magnetic beads. For this, HNRNPC antibody (Santa Cruz sc-32308) or mouse IgG (Jackson 015-000-003) was first purified using Protein A/G beads (Thermo). Briefly, 3 μg of antibody were bound to 15 μl Protein A/G beads (per IP replicate) in Modified Coupling buffer (20 mM sodium phosphate pH 7.2, 315 mM NaCl, 0.1 mM EDTA, 0.1% IGEPAL CA-630, 0.5% glycerol) and incubated 15 min at room temperature. Then the beads were washed twice in Modified Coupling buffer, once in Coupling buffer (20 mM sodium phosphate pH 7.2, 300 mM NaCl) and the antibody eluted in 0.1 M sodium citrate buffer (pH 2.5) for 5 min at room temperature. After neutralization with 1/10 volume of 1 M sodium phosphate buffer (pH 8) the antibody was coupled to M270 Epoxy Dynabeads (Thermo Scientific) in ammonium sulfate buffer (0.1 M sodium phosphate pH 7.4, 1.2 M ammonium sulfate, final concentration) overnight at 37°C. Prior usage, the antibody conjugated beads were washed 4 times in 1X PBS, once in 1X PBS supplemented with 0.5% Tween-20 and resuspended in 1X PBS.

Protein complexes were immunoprecipitated with antibody-conjugated beads for 2h at 4°C, washed three times in wash buffer (15 mM Tris HCl pH 7.5, 150 mM NaCl, 0.1% Triton X-100) and eluted in 1X NuPage LDS sample buffer with 0.1 M DTT for 10 min at 70°C. Eluates were then subjected to alkylation, detergent removal, and Trypsin digestion using Filter Aided Sample Preparation (FASP) protocol^57^, followed by desalting using StageTips^58^. Desalted peptides were subsequently lyophilized by vacuum centrifugation, resuspended in 7 μL of A* buffer (2% ACN, 0.5% Acetic acid, 0.1% TFA in water), and analyzed on a Q-Exactive plus Orbitrap mass spectrometer coupled with a nanoflow ultimate 3000 RSL nano HPLC platform (Thermo Fisher), as described before^59^. Briefly, 6 μL of each peptide sample was resolved at 250 nL/min flow-rate on an Easy-Spray 50 cm x 75 μm RSLC C18 column (Thermo Fisher), using a 123 minutes gradient of 3% to 35% of buffer B (0.1% formic acid in acetonitrile) against buffer A (0.1% formic acid in water), followed by online infusion into the mass spectrometer by electrospray (1.95 kV, 255C). The mass spectrometer was operated in data dependent positive mode. A TOP15 method in which each MS scan is followed by 15 MS/MS scans was applied. The scans were acquired at 375-1500 m/z range, with a resolution of 70,000 (MS) and 17,500 (MS/MS). A 30 seconds dynamic exclusion was applied.

For Co-IP/WB analysis, when indicated, the lysates were pretreated with RNaseA (10 μg RNaseA per 1 mg lysate, 10 min on ice) and incubated with HNRNPC antibody or mouse IgG overnight at +4C. The protein complexes were then immunoprecipitated with Protein A/G beads for 2h at +4C, washed three times with wash buffer (15 mM Tris HCl pH 7.5, 150 mM NaCl, 0.1% Triton X-100) and eluted in 1X NuPage LDS sample buffer with 0.1 M DTT for 10 min at 70C.

### Metastatic colonization assay

Seven-to twelve-week-old age-matched female NOD *scid* gamma mice (NSG, Jackson Labs, 005557) were used for lung colonization assays. For this assay, cancer cells constitutively expressing luciferase were suspended in 100 μL PBS and then injected via tail-vein (2.5×10^4^ for MDA-LM2, 5×10^4^ for MDA-MB-231, 1×10^5^ for HCC1806-LM2). Each cohort contained 4-5 mice, which in NSG background is enough to observe a >2-fold difference with 90% confidence. Mice were randomly assigned into cohorts. Cancer cell growth was monitored *in vivo* at the indicated times by retro-orbital injection of 100 μl of 15 mg/mL luciferin (Perkin Elmer) dissolved in 1X PBS, and then measuring the resulting bioluminescence with an IVIS instrument and Living Image software (Perkin Elmer).

### Histology

For gross macroscopic metastatic nodule visualization, mice lungs (from each cohort) were extracted at the endpoint of each experiment, and 5 μm thick lung tissue sections were hematoxylin and eosin (H&E) stained. The number of macroscopic nodules was then recorded for each section. An unpaired t-test was used to test for significant variations.

### Patient-derived xenografts

Primary tumors of established triple-negative breast cancer PDX models (HCI-001, HCI-002, HCI-010, STG139) were generated in NSG mice as described before^20,21^, collected at a tumor size of 1.0 cm diameter and snap frozen immediately. Tumors were stored at –80ºC until further processed for ribosomal profiling as described above.

### Computational tools

#### Ribolog

Unlike differential gene expression analysis using RNA-seq data which involves comparing two or more count numbers, modeling changes in translational efficiency (TE) requires comparisons of ratios between conditions (TE corresponds to a ratio between ribosome protected mRNA footprint and mRNA abundance counts). The main outcome of interest in ribosome profiling, translation efficiency ratio (TER), is a ratio of two ratios. Since the introduction of ribosome profiling, several analytical packages have been developed that largely inherit the assumptions of prior methods originally designed for RNA sequencing data analysis^60^. A closer evaluation of the underlying assumptions used in many of these tools, e.g. negative binomial (NB) distribution of read counts, revealed that reliable estimation of parameters such as overdispersion required many more biological replicates than are commonly generated in studies of translational control^61,62^. Moreover, the ratio of two NB variables does not follow any known statistical distribution, therefore inference on TER using parametric significance tests remains a challenge. We thus sought to devise a new analytical framework that allows for reliable comparison of TEs across conditions with as few *a priori* assumptions as possible. The resulting method, which we have named Ribolog, relies on logistic regression to model individual Ribo-seq and RNA-seq reads in order to estimate logTER (i.e. log fold-change in TE) and its associated *p*-value across the coding transcriptome.

Before entering the Ribolog pipeline, RNA and RPF fastq files are pre-processed, as described above. Sorted and indexed bam files are imported into Ribolog, and mapped reads are assigned to specific codons using functions borrowed from the R package riboWaltz^63^.

#### Stalling bias detection and correction

Suboptimal codons, RNA secondary structures and activation of RNA binding protein (RBP) binding sites may stall translation at certain codons and produce peaks of RPF reads that stand out against the coding sequence (CDS) background. If not removed, stalling reads will be counted in with other RPF reads and lead to an overestimation of translation efficiency. This may lead to false inference because stalling reads signify locally obstructed translation and should not be misconstrued as a sign of overall increase in translation rate. We developed a new metric, the CELP bias coefficient (CELP: Consistent Excess of Loess Predictions) to measure the strength of stalling bias at each codon. First, we smooth out the observed codon counts along the transcript for each sample using the loess function to produce loess predicted counts. Then, we calculate the CELP bias coefficient and the bias-corrected read count as:

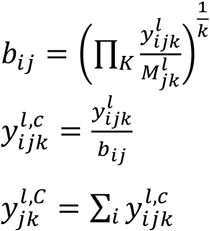

Where:

*b*_*ij*_ : CELP stalling bias coefficient at codon i, gene j

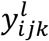: Loess predicted (smoothed) read count at codon i, gene j, sample k

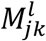: Median of non-zero loess-smoothed counts in gene j, sample k

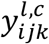: Bias-corrected read count at codon i, gene j, sample k

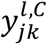: Bias-corrected RPF read count of gene j, sample k

The reasons for using loess-smoothed counts and not raw counts in the above calculations are three-fold: 1) In our experience with multiple ribosome profiling datasets, we have observed that stalling peaks often appear in the same approximate position, but not necessarily the same exact codon, even among replicates of a single biological sample. 2) Some of the factors that impede translation e.g. RBP-binding or RNA secondary structures affect several adjacent codons, not a single codon. 3) Calculation of P-site offset and assignment of RPF reads to specific codons carries a degree of uncertainty, because the distance of read ends from start or stop codon which is used to calculate P-site offset is always a distribution, not a single value, even for reads of the same length. It is therefore beneficial to borrow information from neighboring codons for detection of stalling events. The radius of this neighborhood—which determines the loess “span” parameter—can be changed by the user (default: 5). Median of loess-smoothed non-zero counts 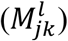 represents background CDS translation level, and the ratio 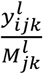shows excess or depletion of reads at any codon position compared with the background i.e. relative peak height. The geometric mean of this ratio among samples produces the bias coefficient for that position. If the goal of the study is to investigate local patterns of stalling between groups of samples, group-specific bias coefficients should be calculated. CELP coefficients can be then regressed against any position-specific (e.g. RBP binding site or codon type), transcript-specific (e.g. length or existence of known upstream ORFs) or group-specific (e.g. wild type vs. tRNA knockdown cell line) factors to infer their effects on stalling. On the other hand, if CELP is primarily used to debias RPF counts to allow an unbiased TER test, all samples in the dataset can be pooled together in the calculation of bias coefficients.

### TER test

Both RNA and RPF libraries are mapped to the same reference transcriptome as described in the previous section. RNA read counts per transcript are calculated directly from the bam files. RPF read counts are either obtained in the same way or run through CELP debiasing first to smooth out local non-uniformities (described above in detail). RNA and RPF transcript x sample count matrices are normalized separately for library size variation using the median-of-ratios method^64^. We model translation efficiency (TE) as the odds of retrieving two different sequencing read types from a sample: RPF vs. RNA. In this scenario, we hypothetically pool all the reads from an experiment, and then extract a read from this pool. The odds of extracting an RPF vs. an RNA read from this pool yields a probabilistic estimation of translation efficiency. We compare translation efficiency ratio (TER) by testing the effect of model covariates on TE, i.e. the odds ratio of RPF/RNA between groups or per unit change of continuous predictors. In the very simple case of comparing only two non-replicated samples, a significance test on TER could be performed using a Chi-square or Fisher’s exact test on a 2×2 contingency table with sample name acting as the exposure (independent variable) and the read type (RPF or RNA) as the response (dependent variable). Since most biological experiments are replicated and involve multiple sample groups, we generalize the test in a logistic regression setting:

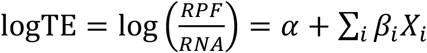

Where:

*RPF*: normalized (and optionally debiased) RPF read count

*RNA*: normalized RNA read count

*α*: intercept

*X*_*i*_: Predictor (independent variable) i

*β*_*i*_: Regression coefficient for predictor (independent variable) i

The test is run separately for each transcript. Independent variables could be categorical e.g. group labels, or continuous to represent a molecular measurement from the sample e.g. tRNA concentrations or a codon optimality score. This formulation of TER accommodates complex experimental designs with any number of groups or replicates described by any number of attributes (covariates). A *p*-value is reported for each regression coefficient indicating the significance of its effect (“effect” here is defined as a regression coefficient being different from 0, or the corresponding TER being different from 1). The effect sizes (logTER) and log10(*p*-values) are plotted together to produce the familiar volcano plot. The expected TER of a transcript between two sample differing in one or multiple attributes can be estimated by substituting the obtained regression coefficients in the equation below:

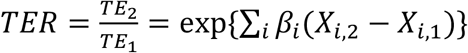

For detailed instructions to install the package, prepare the input data, run the tests, and interpret and plot the results, visit https://github.com/goodarzilab/Ribolog. Additional modules for quality control (QC), empirical null significance testing to reduce false positives, meta-analysis of ribosome profiling data, etc. are available from the github page.

### APAlog

The number of RNA reads mapping to each poly(A) site of a transcript could originate from a regular RNA-seq or a specialized 3’ UTR sequencing protocol. Normalized counts are then used by APAlog to assess the extent and pattern of differential poly(A) site usage via multinomial logistic regression:

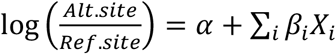

Where:

*Alt. site*: Alternative poly(A) site normalized read count

*Ref. site*: Reference poly(A) site normalized read count

*α*: intercept

*X*_*i*_: Predictor (independent variable) i

*β*_*i*_: Regression coefficient for predictor (independent variable) i

APAlog automatically sets the poly(A) site of each transcript that comes first alphabetically to reference. The user can specify which poly(A) site to serve as reference by adjusting the poly(A) site names in the count matrix. APAlog can be run in three modes: 1) Overall transcript-wise test: A deviance test is performed between the fitted model with covariates and the null (intercept-only) model. This test identifies transcripts which show differential poly(A) site selection among samples but does not specify which poly(A) sites or covariates contribute to the difference. This mode facilitates the quick scanning of a large multi-group dataset to flag putative targets of regulation. Moreover, performing exactly one test per transcript, it avoids complications of multiple testing correction among transcripts with unequal number of poly(A) sites. 2) Alternatives vs. reference test: One poly(A) site per transcript is marked as the reference site and all others (one or more) are tested against it. This mode is suitable for specific applications such as testing 3’ UTR length variation when one poly(A) site, in this case the most proximal one, can be set to reference and all others compared to it. 3) Pairwise test: This test compares all pairs of poly(A) sites per transcript and provides the highest resolution view of poly(A) site selection regulation. It is also the best choice if a reference or canonical poly(A) site cannot be logically assigned.

For detailed instructions to install the package, prepare the input data, run the tests and interpret the results, visit https://github.com/goodarzilab/APAlog.

## ACKNOWLEDGEMENTS

We acknowledge the UCSF Center for Advanced Technology (CAT) for high throughput sequencing and other genomic analyses. We thank S. F. Tavazoie for the gift of the HCC1806 and HCC1806-LM2 cell lines. We thank the Preclinical Therapeutics core as well as the Laboratory Animal Resource Center (LARC) at UCSF. We acknowledge support from our colleagues at the Helen Diller Family Comprehensive Cancer Center and the Breast Oncology Program.

## Funding

This work was supported by grants from the NIH (R00CA194077 and R01CA240984) and ACS (130920-RSG-17-114-01-RMC) to H.G. This research was also supported by funding from the UCSF Helen Diller Family Comprehensive Cancer Center Breast Oncology Program (the content is solely the responsibility of the authors). This study was supported in part by UCSF Laboratory for Cell Analysis shared resource facility through a grant from NIH (P30CA082103). This work used the Vincent J. Proteomics/Mass Spectrometry Laboratory at UC Berkeley, supported in part by NIH S10 Instrumentation Grant S10RR025622. L.F. was supported by NIH training grant T32CA108462-15. A.N. was supported by DoD PRCRP Horizon Award W81XWH-19-1-0594. S.Z. was supported by an HHMI medical research fellowship. M.D. and F.K.M. were supported by an MRC career development award to F.K.M. (MR/P009417/1). D.M. was supported by an MD fellowship from the Boehringer Ingelheim Fonds. J.W. was supported by an EMBO long-term post-doctoral fellowship (EMBO ALTF 159-2017). A.G. and J.W. were supported by U01 CA199315, Mark Foundation and CDMRP DoD Breakthrough Award W81XWH-16-1-0603. In the end, we acknowledge the support of our late colleague Dr. Zena Werb for the PDX studies reported here.

## AUTHOR CONTRIBUTIONS

H.G. and A.G. conceptualized the study. A.N. performed Ribo-seq, RNA-seq, PAPERCLIP, CoIP, TMT labeling, western blotting, qPCR experiments. L.F. performed CLIP-seq. J.W. performed PDX transplantation experiments. H.A. and S.M. developed Ribolog and APAlog. A.N., K.G., D.M., B.C., S.Z., T.J., P.N., N.S. generated cell lines and performed *in vivo* metastasis experiments. A.N. and P.N. performed T4 treatment experiments. M.D. and F.K.M. performed mass spectrometry and analysis. H.W.H. contributed to mouse experiments. H.G. analyzed the Ribo-seq, RNA-seq data, TCGA data, and clinical data. A.N., H.A., L.F., A.G, and H.G. wrote the manuscript. H.G. supervised all research.

## COMPETING INTERESTS STATEMENT

Authors declare no competing interests.

## FIGURE LEGENDS

**Supplementary Figure 1.**
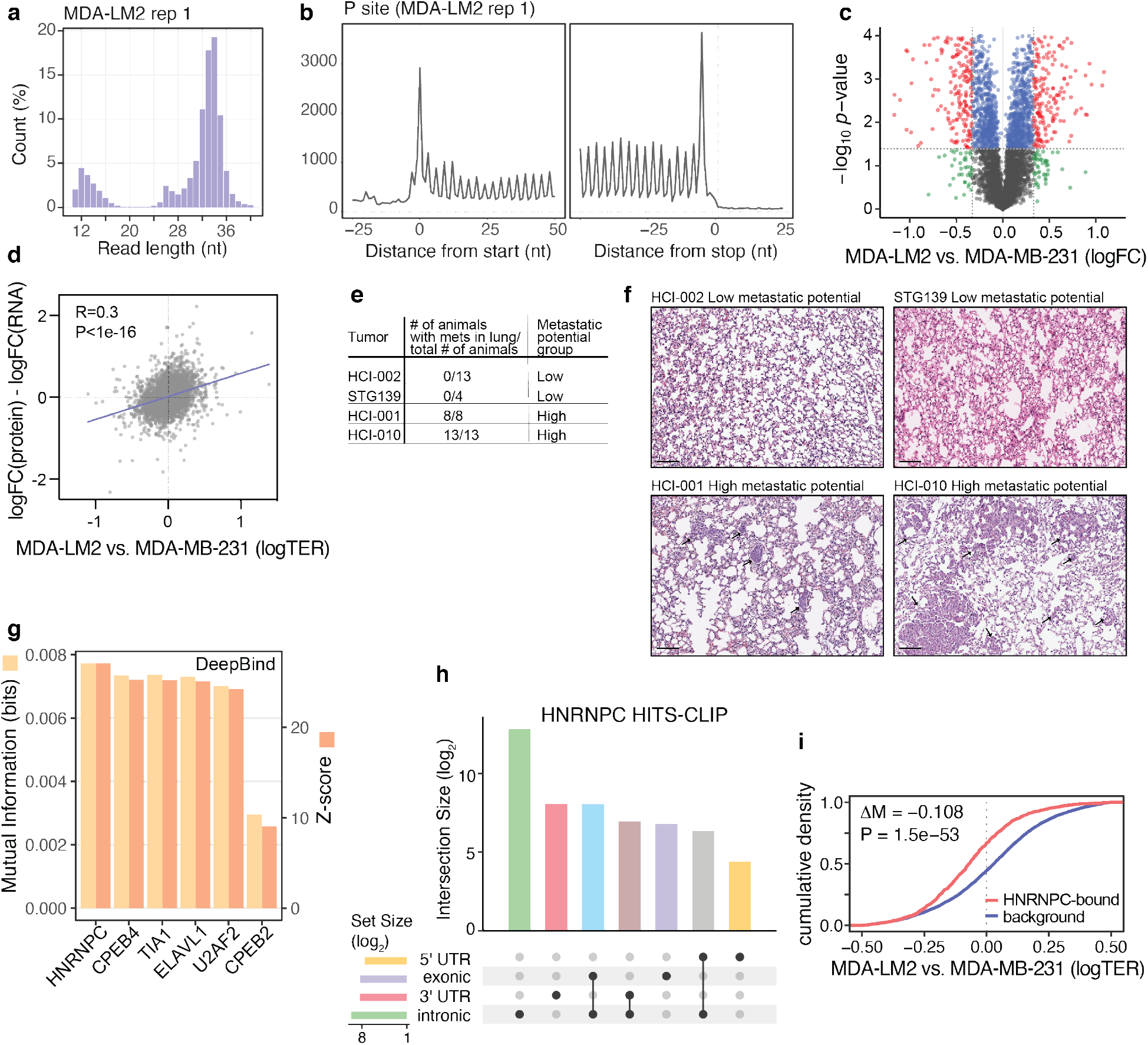
**(a)** Length distribution of ribosome protected footprints (RPFs) as determined by Ribo-seq. A representative data sample (MDA-LM2 cells, replicate 1) is shown. **(b)** Distribution of RPFs, aligned on an inferred ribosome P-site, on a metagene, centered around translation start (left) or stop (right) site. A representative data sample (MDA-LM2 cells, replicate 1) is shown. **(c)** Volcano plot illustrating the changes in protein abundance in MDA-LM2 compared to MDA-MB-231 cells, as determined by TMT-MS analysis. The data points are colored according to thresholds in effect size (logFC ± 0.33) and significance (*p* < 0.05, *t*-test). **(d)** The distribution of changes in TEs (as determined by Ribo-seq) and in protein abundance (as determined by TMT-MS and normalized by RNA expression obtained from RNA-seq), in MDA-LM2 compared to MDA-MB-231 cells. Pearson R and associated *p* value are shown. **(e)** The comparison of the metastatic capacity of breast cancer PDXs used in this study. **(f)** Representative images of H&E stained mouse lung sections transplanted with breast cancer PDXs. The metastatic foci are indicated by black arrows. **(g)** Mutual information (MI) values and associated *z-*scores from the DeepBind algorithm, showing the prediction of poly(U) binding protein targets among translationally repressed mRNAs in MDA-LM2 compared to MDA-MB-231 cells. **(h)** Upset plot showing the distribution and overlap of HNRNPC peaks within genomic features, as determined by CLIP-seq. **(i)** Cumulative density plot of translation efficiency ratios (TER) comparing MDA-LM2 to MDA-MB-231 cells, for HNRNPC 3’ UTR target and non-target mRNAs. Median difference (ΔM) and *p* value (calculated using Mann-Whitney *U*-test) are shown.

**Supplementary Figure 2.**
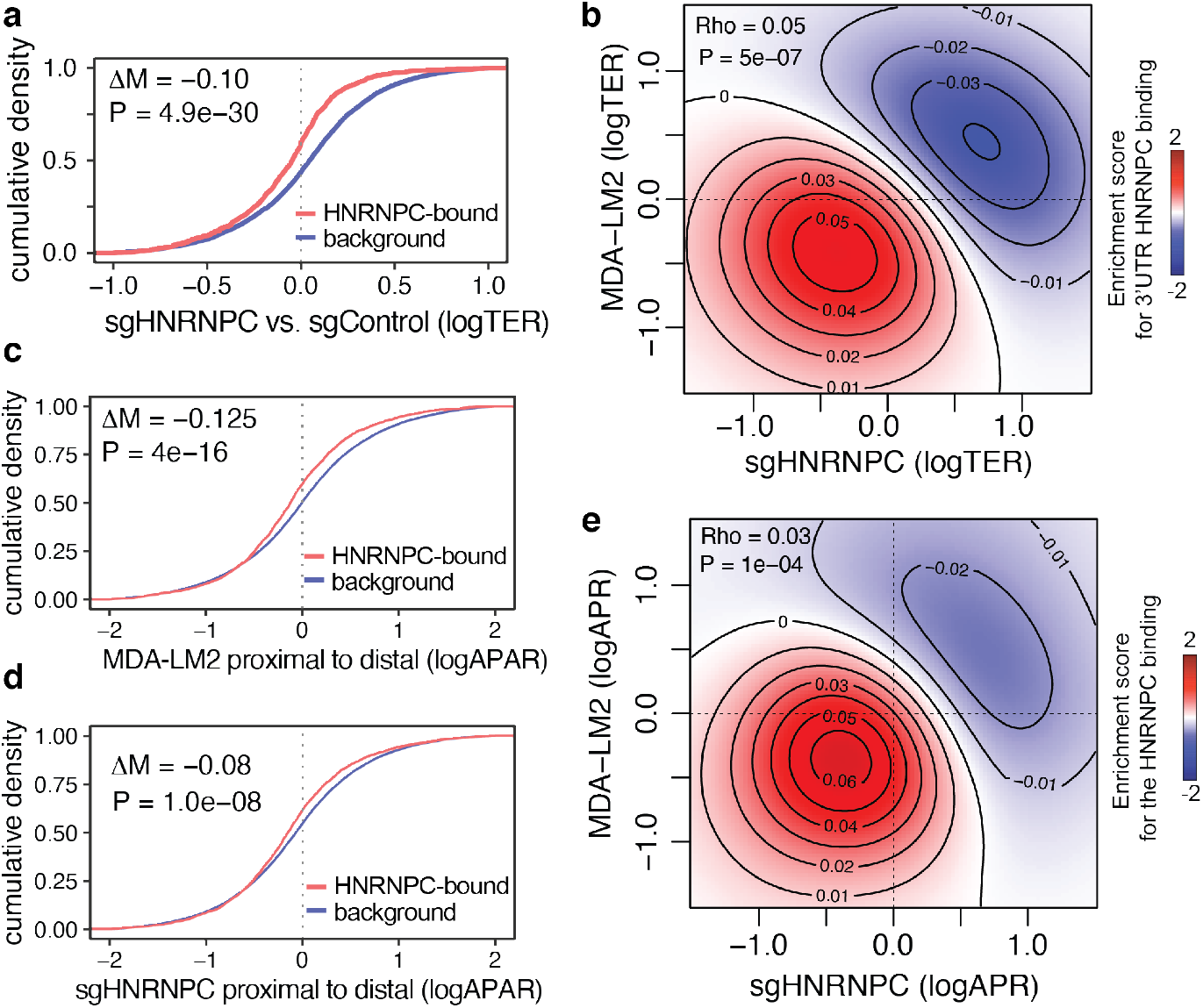
**(a)** Cumulative density plot of translation efficiency ratios (TER) comparing sgHNRNPC to sgControl MDA-MB-231 cells, for HNRNPC 3’ UTR target and non-target mRNAs. Median difference (ΔM) and *p* value (calculated using Mann-Whitney *U*-test) are shown. **(b)** Two-dimensional heatmap showing significant logTER correlation of translationally repressed mRNAs in MDA-LM2 and HNRNPC knockdown (sgHNRNPC) cells. For comparison, the Spearman correlation coefficient and the associated *p* value are shown across all genes. **(c)** Cumulative density plot of alternative polyadenylation ratios (logAPAR) comparing MDA-LM2 to MDA-MB-231 cells, for HNRNPC 3’ UTR target and non-target mRNAs; statistics as in (a). **(d)** Cumulative density plot of logAPAR comparing sgHNRNPC to sgControl cells, for HNRNPC 3’ UTR target and non-target mRNAs; statistics as in (a). **(e)** Two-dimensional heatmap showing significant logAPAR correlation of proximal to distal poly(A) site switch in MDA-LM2 and HNRNPC knockdown (sgHNRNPC) cells; statistics as in (b).

**Supplementary Figure 3.**
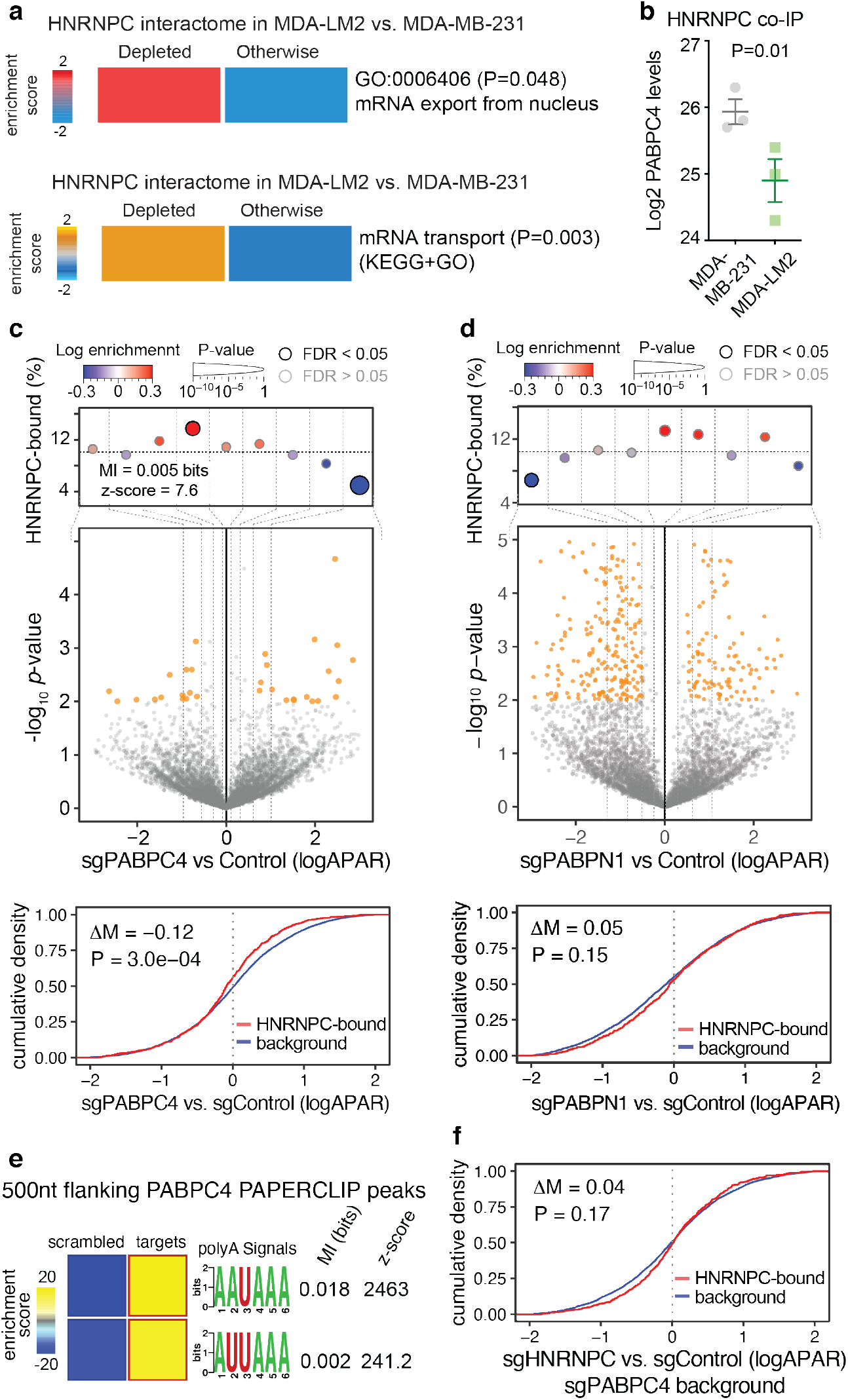
**(a)** Significant depletion of selected gene ontology (GO) terms in the HNRNPC interactome in MDA-LM2 compared to MDA-MB-231 cells, as determined by coIP-MS. Also reported are the associated empirical *p* values. **(b)** Enrichment of PABPC4 in HNRNPC coIP-MS data in MDA-MB-231 and MDA-LM2 cells. *P* value calculated using *t*-test. **(c)** Middle: Volcano plot showing distribution of changes in alternative polyadenylation ratio (logAPAR) in sgPABPC4 compared to sgControl MDA-MB-231 cells. Statistically significant (logistic regression, *p* < 0.01) observations are highlighted in orange. Top: Enrichment of the HNRNPC targets as a function of logAPAR between sgHNRNPC/sgPABPC4 and sgControl/sgPABPC4 cells. mRNAs are distributed to equally populated bins according to their logAPAR (dotted vertical lines delineate the bins); the y-axis shows the frequency of the HNRNPC-bound 3’ UTRs that we identified in each bin (dotted horizontal line denotes the average HNRNPC target frequency across all transcripts). Bins with significant enrichment (logistic regression, FDR < 0.05; red) or depletion (blue) of HNRNPC targets are denoted with a black border. Also included are mutual information (MI) values and their associated *z*-scores. Bottom: Cumulative density plot of logAPAR comparing sgPABPC4 to sgControl cells, for HNRNPC 3’ UTR target and non-target mRNAs. Median difference (ΔM) and *p* value (calculated using Mann-Whitney *U-*test) are shown. **(d)** Comparison of logAPAR in sgPABPN1 and sgControl cells, as in (c). **(e)** Heatmaps showing the enrichment of canonical poly(A) signals in the vicinity of PABPC4 binding peaks (as determined by PAPER-CLIP). The bolded border denotes a statistically significant enrichment (hypergeometric test, corrected *p* < 0.05; red). MI values and associated *z*-scores are shown. **(f)** Cumulative density plot of logAPAR comparing sgHNRNPC/sgPABPC4 (double knockdown) to sgControl/sgPABPC4 cells, for HNRNPC 3’ UTR target and non-target mRNAs; statistics as in (c).

**Supplementary Figure 4.**
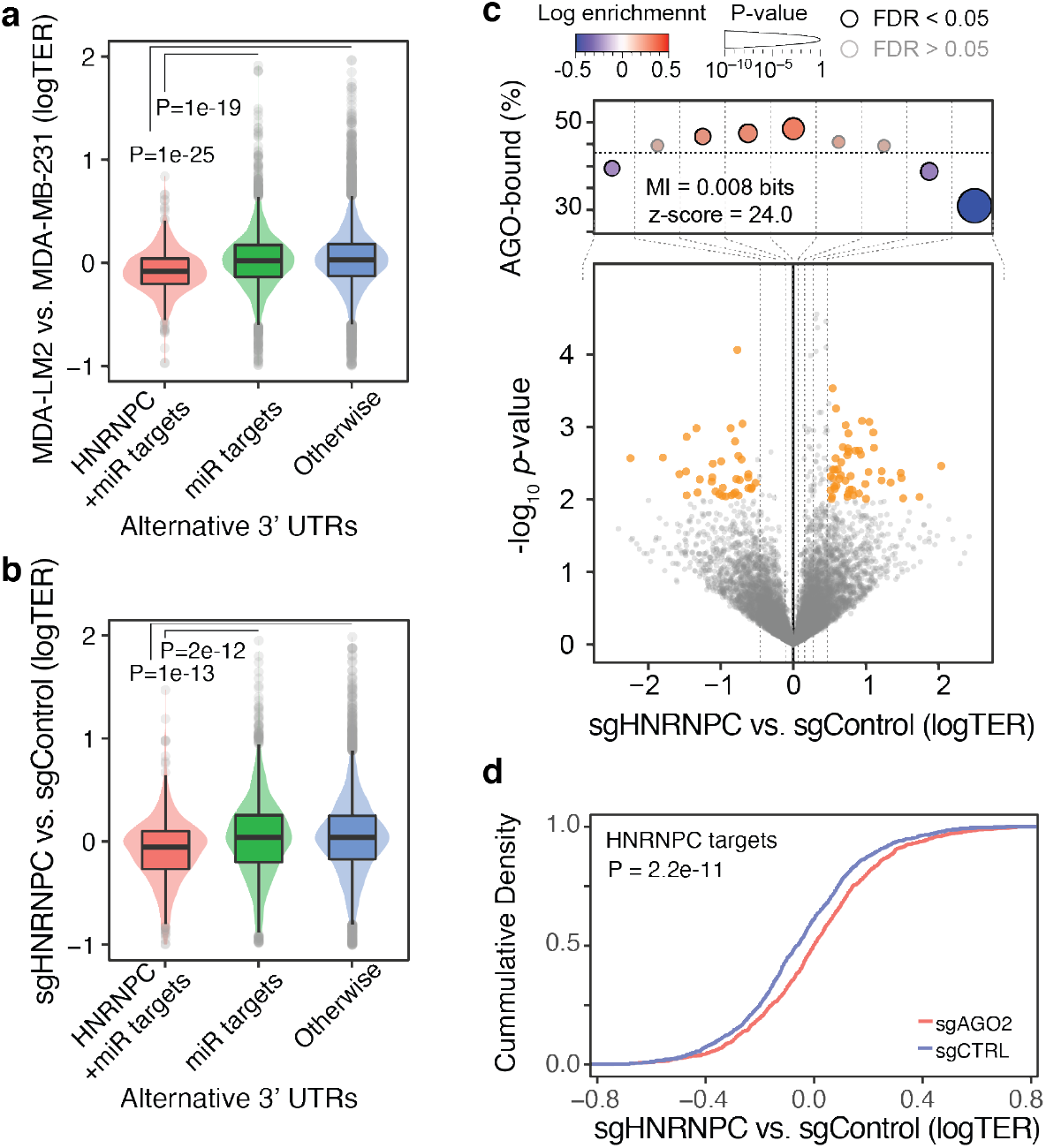
**(a)** Violin plots showing the distribution of translation efficiency ratios (logTER) comparing MDA-LM2 to MDA-MB-231 cells among the miRNA target, joint HNRNPC and miRNA target, and non-target mRNA 3’ UTRs. *P* values calculated using Mann-Whitney *U*-test. **(b)** Violin plots showing the distribution of translation efficiency ratios (logTER) comparing sgHNRNPC to sgControl cells among the miRNA target, joint HNRNPC and miRNA target, and non-target mRNA 3’ UTRs. *P* values calculated using Mann-Whitney *U*-test. **(c)** Bottom: Volcano plot showing distribution of changes in translation efficiency ratio (logTER) in sgHNRNPC compared to sgControl cells. Top: Enrichment of the AGO2 targets as a function of logTER between sgHNRNPC and sgControl cells; statistics as in Fig. 2a. **(d)** Cumulative density plot of logTER (HNRNPC 3’ UTR targets) comparing sgHNRNPC to sgControl cells, in AGO2 knockdown (sgAGO2) and control (sgControl) conditions; statistics as in Fig. S2a.

**Supplementary Figure 5.**
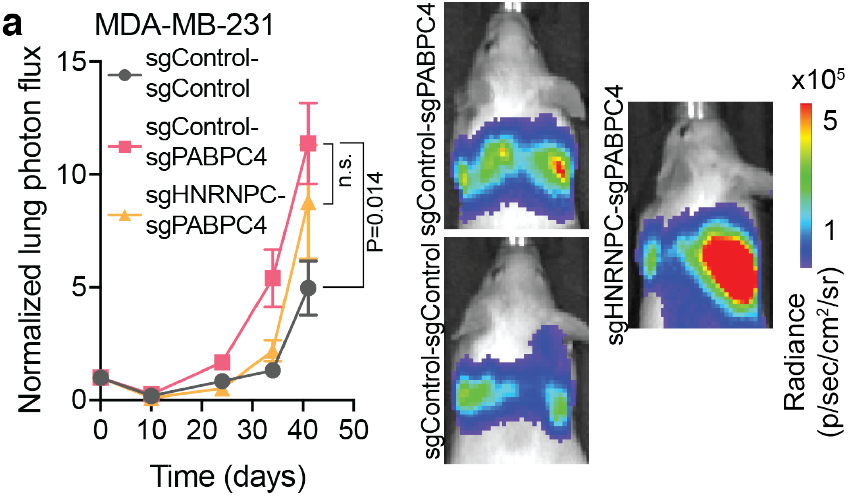
**(a)** MDA-MB-231 cells stably expressing sgControl, sgPABPC4 or sgPABPC4/sgHNRNPC were injected via tail vein into NSG mice. Bioluminescence was measured at the indicated times (*p* value calculated using two-way ANOVA). *n =* 4-5 mice per cohort.

**Supplementary figure 6.**
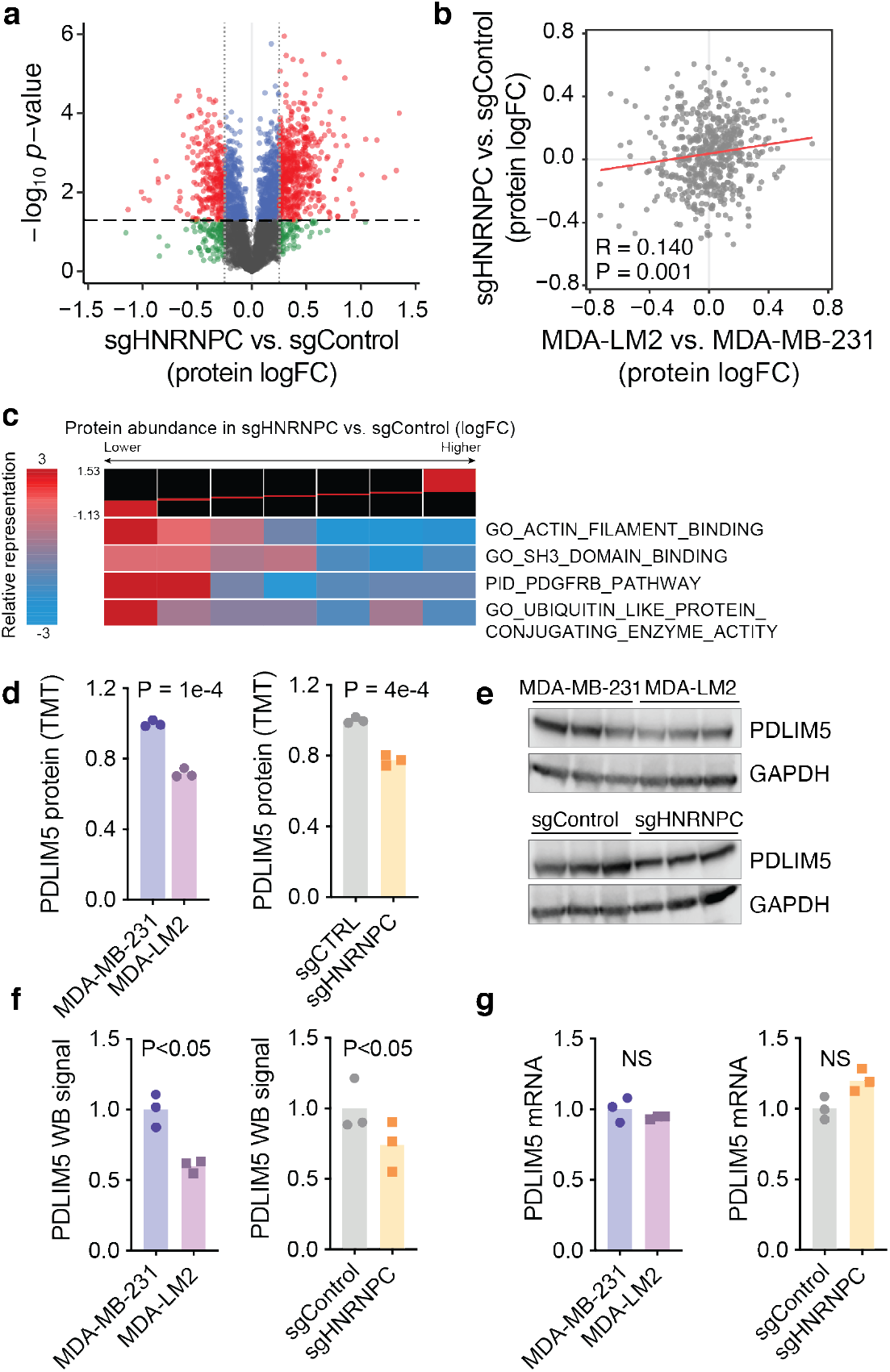
**(a)** Volcano plot illustrating the changes in protein abundance in HNRNPC knockdown (sgHNRNPC) compared to control (sgControl) MDA-MB-231 cells, as determined by TMT-MS analysis. The data points are colored according to thresholds in effect size (logFC *±* 0.25) and significance (*p* < 0.05, *t-*test). **(b)** The distribution of changes in protein abundance in MDA-LM2 vs. MDA-MB-231 cells and sgHNRNPC vs. sgControl cells, as determined by TMT-MS. Pearson R and associated *p* value are shown. **(c)** Gene-set enrichment analysis of the data depicted in (a). The proteins were distributed into equally populated based on their logFC in sgHNRNPC vs. sgControl cells. Then the enrichment of a given gene set was calculated in each bin using iPAGE^19^, a mutual information-based algorithm. **(d)** Quantification of PDLIM5 protein expression in MDA-MB-231 and MDA-LM2 (left), or sgControl and sgHNRNPC (right) cells, as determined by TMT-MS. *P* values calculated using one-tailed Student’s *t*-test. **(e)** Western blot analysis of PDLIM5 and GAPDH (loading control) expression in MDA-MB-231 and MDA-LM2 (top), or sgControl and sgHNRNPC (bottom) cells. **(e)** Quantification of relative PDLIM5 protein expression (normalized to GAPDH) in MDA-MB-231 and MDA-LM2 cells (left) or sgControl and sgHNRNPC cells (right), as determined by western blotting in (c). *P* values calculated using one-tailed Mann-Whitney *U*-test. **(f)** Quantification of relative PDLIM5 mRNA expression (normalized to HPRT) in MDA-MB-231 and MDA-LM2 cells (left) or sgControl and sgHNRNPC cells (right), as determined by RTqPCR.

**Supplementary Figure 7.**
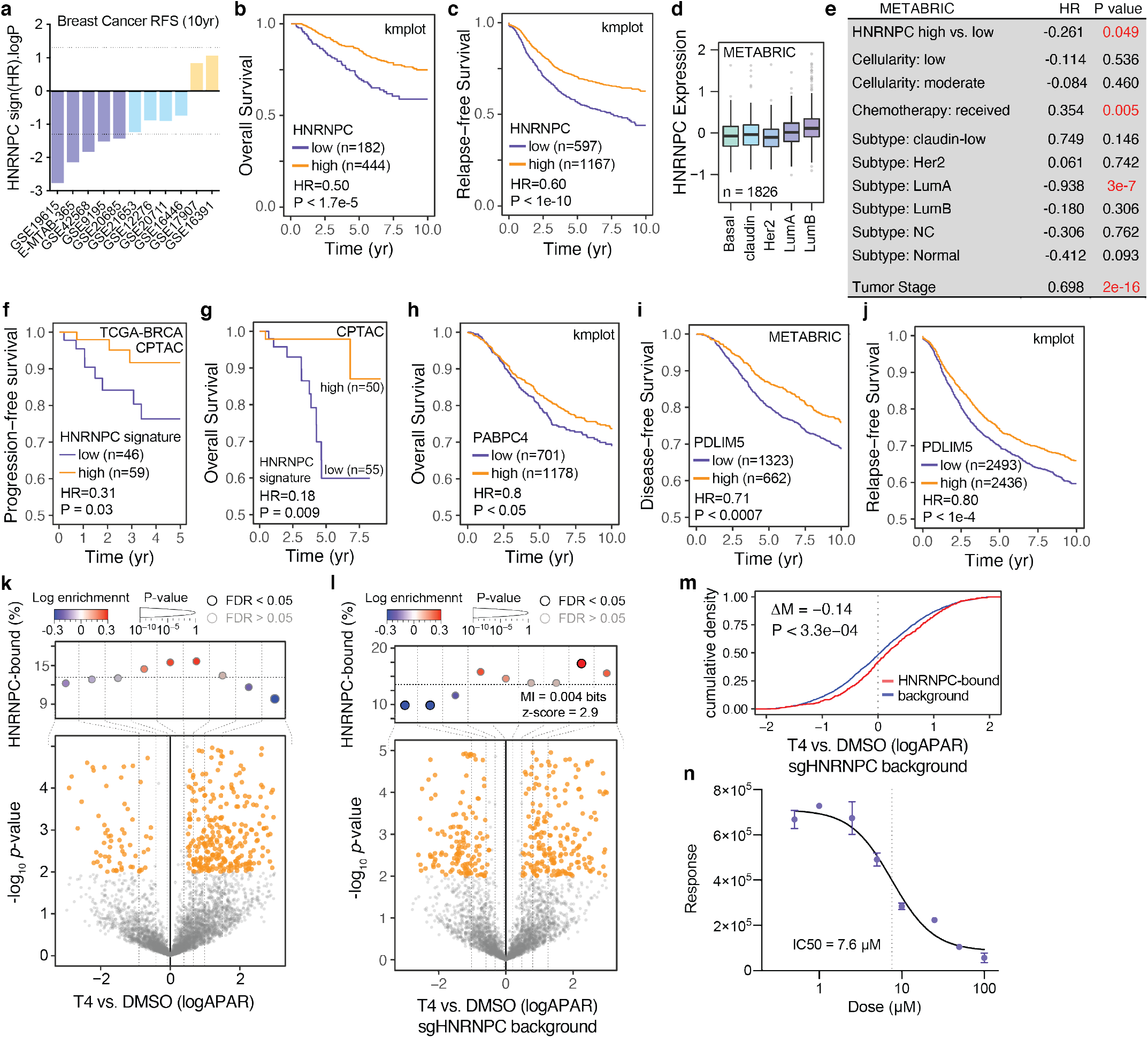
**(a)** Shown are distribution of 10-year relapse-free survival *p* values (two-sided log rank test results reported as –log *p* for positive association and log *p* for negative) of the association of HNRNPC expression and clinical outcome in the listed 10 breast cancer datasets. Violet bars show associations that pass the statistical threshold (–log *p* < –1.3, FDR-corrected two-sided log-rank test (FDR < 0.1)), blue bars are trending negative, and yellow bars are trending positive. The statistical threshold was adjusted as 10/number of datasets. **(b)** Kaplan-Meier survival curve showing association between tumor HNRNPC levels and overall survival in a collection of breast cancer patient cohorts. **(c)** Kaplan-Meier survival curve showing association between tumor HNRNPC levels and relapse-free survival in a collection of breast cancer patient cohorts. **(d)** HNRNPC mRNA levels across breast tumor subtypes in the METABRIC cohort. **(e)** Multivariate survival analysis (Cox proportionate-hazards model) of breast cancer patients in the METABRIC cohort with HNRNPC expression as one of the factors. *P* < 0.05 are highlighted in red. LumA, luminal A; LumB, luminal B; NC, not classified. **(f)** Kaplan-Meier survival curve showing association between tumor HNRNPC signature protein levels and progression-free survival in the TCGA-BRCA CPTAC cohort. **(g)** Kaplan-Meier survival curve showing association between tumor HNRNPC signature protein levels and overall survival in the TCGA-BRCA CPTAC cohort. **(h)** Kaplan-Meier survival curve showing association between tumor PABPC4 levels and overall survival in a collection of breast cancer patient cohorts. **(i)** Kaplan-Meier survival curve showing association between tumor PDLIM5 expression and disease-free survival in the METABRIC cohort. **(j)** Kaplan-Meier survival curve showing association between tumor PDLIM5 levels and relapse-free survival in a collection of breast cancer patient cohorts. Hazard ratios (HR) and *p* values (calculated using log-rank test) are shown. **(k)** Bottom: Volcano plot showing distribution of changes in alternative polyadenylation ratio (logAPAR) in T4-compared to vehicle control (DMSO)-treated MDA-MB-231 cells. Top: Enrichment of the HNRNPC targets as a function of logAPAR between T4-and DMSO-treated cells; statistics as in Fig. S3c. **(l)** Bottom: Volcano plot showing distribution of changes in logAPAR in T4-compared to vehicle control (DMSO)-treated HNRNPC knockdown (sgHNRNPC) cells. Top: Enrichment of the HNRNPC targets as a function of logAPAR between T4- and DMSO-treated cells; statistics as in Fig. S3c. **(m)** Cumulative density plot of logAPAR comparing T4- to DMSO-treated HNRNPC knockdown (sgHNRNPC) cells; statistics as in Fig. S2a. **(n)** Dose-response measurements for 6-hour T4 treatment and corresponding cell viability, determined 72 hours post-treatment.

## Notes

### Competing Interest Statement

The authors have declared no competing interest.

